# Deep dynamical modelling of developmental trajectories with temporal transcriptomics

**DOI:** 10.1101/2023.07.06.547989

**Authors:** Rory J. Maizels, Daniel M. Snell, James Briscoe

## Abstract

Developmental cell fate decisions are dynamic processes driven by the complex behaviour of gene regulatory networks. A challenge in studying these processes using single-cell genomics is that the data provides only a static snapshot with no detail of dynamics. Metabolic labelling and splicing can provide time-resolved information, but current methods have limitations. Here, we present experimental and computational methods that overcome these limitations to allow dynamical modelling of gene expression from single-cell data. We developed sci-FATE2, an optimised metabolic labelling method that substantially increases data quality, and profiled approximately 45,000 embryonic stem cells differentiating into multiple neural tube identities. To recover dynamics, we developed velvet, a deep learning framework that extends beyond instantaneous velocity estimation by modelling gene expression dynamics through a neural stochastic differential equation system within a variational autoencoder. Velvet outperforms current velocity tools across quantitative benchmarks, and predicts trajectory distributions that accurately recapitulate underlying dataset distributions while conserving known biology. Velvet trajectory distributions capture dynamical aspects such as decision boundaries between alternative fates and correlative gene regulatory structure. Using velvet to provide a dynamical description of in vitro neural patterning, we highlight a process of sequential decision making and fate-specific patterns of developmental signalling. Together, these experimental and computational methods recast single-cell analyses from descriptions of observed data distributions to models of the dynamics that generated them, providing a new framework for investigating developmental gene regulation and cell fate decisions.

## INTRODUCTION

The formation of patterned, functional tissues depends on organised cell fate determination. This is driven by gene regulatory networks (GRNs) that interpret extra-cellular signals to induce the correct cellular responses [1]. Single-cell RNA sequencing studies have illustrated the complexity of these developmental gene regulatory networks, documenting, for example, the temporal patterns of transcription factor expression in the developing nervous system [2] and defining sequences of cellular states through development [3, 4].

Progressing beyond descriptive analyses of heterogeneous phenotypes to modelling the underlying regulatory mechanisms has long been a major goal in the single-cell field [5], but remains a difficult task. Gene regulatory networks are complex and time-dependent, with feedback cycles and emergent behaviours. To understand such complex processes, dynamical modelling is required [6].

However, single-cell sequencing provides only a static snapshot of cellular composition. Dynamics must be inferred through pseudo-temporal ordering [7], which collapses the tens of thousands of data points commonly found in single-cell experiments into a single, aggregated time series. This can provide valuable descriptive insight [8] but lacks the resolution to capture causal relationships between genes (for example, pseudotime information does not benefit GRN inference algorithms [9, 10]). Sufficiently capturing the complex dynamics of developmental regulation will be necessary to move from descriptive analyses to quantitative models of causal mechanisms; an important step for this is to capture temporal dynamics at single-cell resolution.

Approaches that resolve time in single cells have been developed: RNA velocity [11] infers dynamics from the ratio of spliced and unspliced reads. This approach has been widely applied [12–16] but it is limited by the technical constraints and confounding factors that arise from relying on gene splicing for temporal information [17–19].

Metabolic labelling of RNA with labels such as thiouridine (4sU) provides a more direct, experimentally controllable measurement of time [20– In this approach, 4sU incorporates into nascent RNA, and is subsequently converted to a cytosine analogue with iodoacetamide (IAA) treatment, allowing new RNA to be distinguished from old based on U-to-C mutations in the sequencing data. Of available methods, only sci-FATE [20], scNT-seq [22] and the more recent Well-TEMP-seq [24] provide throughput comparable to commercial single-cell platforms, and only sci-FATE can do so without the use of custom microfluidics or microwell devices. However, sci-FATE, based on combinatorial indexing, is a labour-intensive and potentially error-prone method, and the method’s cell treatment protocol is highly destructive, with treatments include liquid nitrogen freezing, PFA, Triton-X, hydrochloric acid and 50% DMSO. As such, initial pilots of sci-FATE revealed low quality data that would heavily restrict the method’s ability to capture complex or subtle regulatory dynamics.

Alongside experimental methods, a number of computational tools have been developed for inference of gene expression ‘velocity’ from either splicing or labelling data [11, 18, 25–30]. In particular, numerous recent studies have applied methods from deep generative modelling [26, 29–31] to the task of velocity inference. By applying the flexibility and power of neural networks, deep generative modelling is particularly well suited to learning hidden variables that capture the complex distributions within high dimensional data. Similar approaches have already been successful in single-cell genomics tasks such as multimodal integration [31], perturbation prediction [32] and data correction [33]. However, issues remain: first, recent work has found that aspects common across velocity analyses such as data smoothing, UMAP embedding and velocity projection methods, can affect the inferred dynamics and lead to inconsistent inferences [17–19]. Second, velocity inference produces estimates of instantaneous velocity describing deterministic cell dynamics, which may be insufficient to accurately capture the stochastic global dynamics of complex developmental systems. By transforming single-cell analyses from descriptions of observed data distributions into models of the dynamics that generated them, time-resolved methods could facilitate a deeper understanding of underlying mechanisms. To achieve this, both experimental and computational challenges must be addressed.

Here we present an integrated framework of experimental and computational methods for modelling stochastic dynamical systems of gene expression using time-resolved single-cell RNA sequencing data. Experimentally, we developed sci-FATE2, an optimised, semi-automated version of sci-FATE [20], providing improvements that result in data quality and cellular throughput comparable to commercial platforms.

Computationally, we developed velvet, a variational autoencoder [34] to model velocity dynamics with a lower-dimensional vector field, constraining predicted velocities based on local neighbourhood information. To extend our analyses beyond estimation of instantaneous velocities, we developed an extended model, velvetSDE, that infers global dynamics by embedding the learnt vector field in a neural stochastic differential equation (nSDE) system [35, 36] that is trained to produce accurate trajectories that stay within the data distribution.

We show through quantitative benchmarking that velvet outperforms existing velocity tools across all metrics, without the need for data smoothing. velvetSDE’s predicted trajectory distributions map the commitment of cells to specific fates over time, and can faithfully conserve known trends while capturing correlative structures between related genes that are not observed in unrelated genes. We use our methods to study neural tube patterning, a complex series of developmental decisions that can be recapitulated in vitro [37, 38]. We collect 44,713 time-resolved transcriptomes, capturing the differentiation of mouse ES cells to neuromesodermal progenitors (NMPs) that transition to neural and mesodermal identities, and the subsequent specification of neural cell types, floor plate, motor neurons and V3 interneurons. Applying our framework to study in vitro neural patterning decisions, we describe how this process occurs as two distinct fate decisions, and resolve differences in the expression of Shh regulators that distinguish different neural fates. Our analysis suggests a degree of variability in signal interpretation and points towards possible strategies for more precise fate manipulation in vitro.

## RESULTS

### Limitations of current velocity methods

RNA velocity requires that both spliced and unspliced mRNA species can be detected for each gene. However, splicing varies significantly across genes. Many transcription factors contain no introns to be spliced. Examining datasets from several tissues from both mouse and human [2, 11–13, 39, 40], we see a substantial proportion of genes with minimal unspliced reads, and this proportion increases for transcription factors and highly variable genes (Figure 1A), such that often over a quarter of genes retained for analysis will have less than 1 in 20 reads being unspliced. Given that generally far fewer than 20 reads are detected per gene per cell (for example, the mean non-zero value across the above datasets is 2.4), this suggests that for many genes, splicing does not provide a robust measurement of nascent transcription at single-cell level. Moreover, it has been suggested that the methods used to project velocity into low-dimensional visualisations may confound assessment of high dimensional velocity vectors [18, 19]. Because it is not possible to directly project velocity vectors into commonly used embeddings such as UMAP and t-SNE, a projection heuristic based on nearest neighbours is used. These projections usually appear to qualitatively capture known biology [12–16], but they introduce a confounding factor when attempting to evaluate the quality of inferred velocities in the original data space. However, velocities can be directly projected into PCA space, allowing a more direct, heuristic-free inspection of high dimensional velocities. Doing so with an example dataset (with velocities inferred using scVelo) revealed substantial differences in the predicted dynamics derived with and without the use of neighbour based transition probabilities (Figure 1B). Directly projected velocities displayed less plausible dynamics, pointing in directions that are not plausible given the distribution of cells in the dataset.

**Figure 1:**
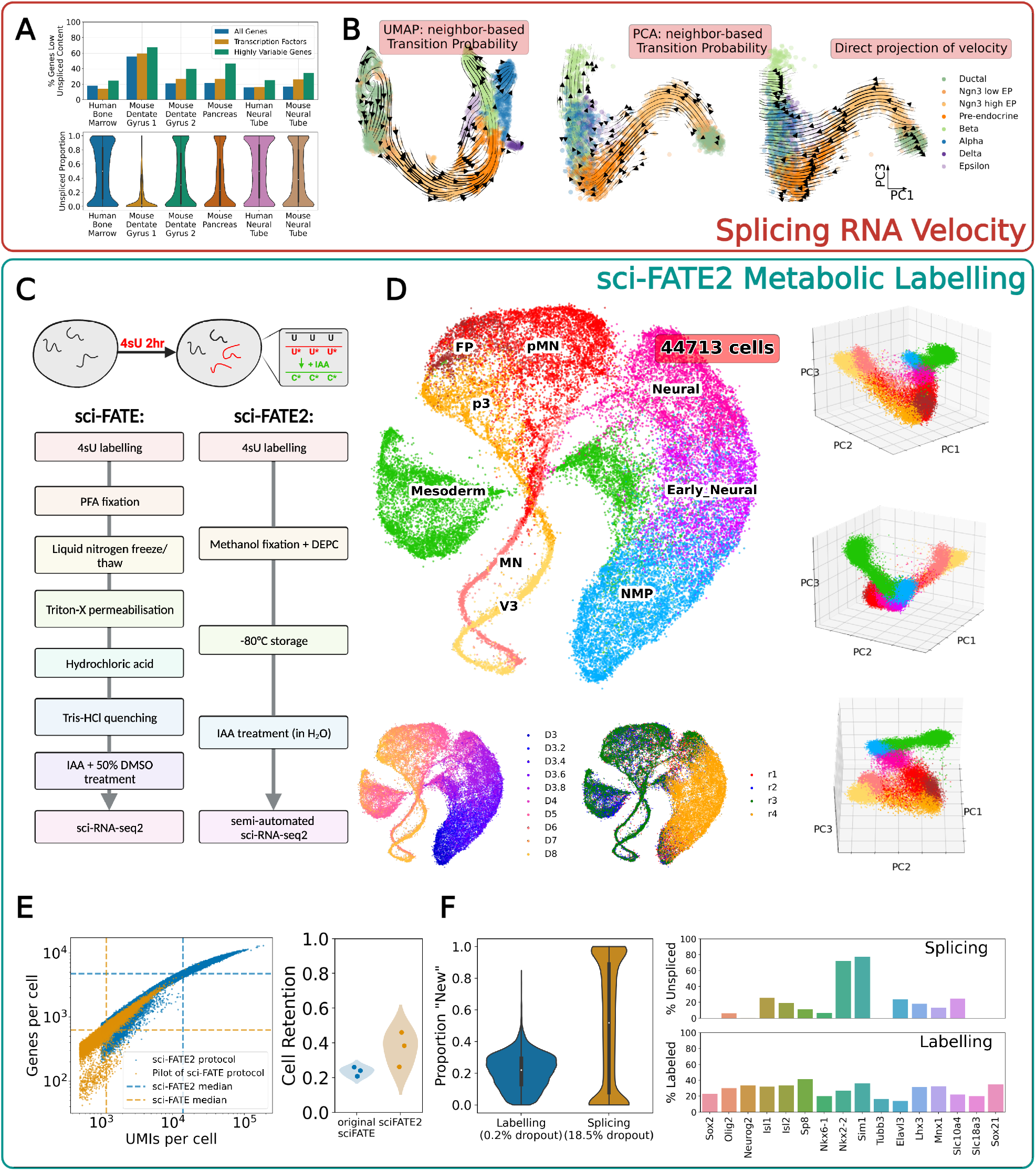
Improved temporal transcriptomics with sci-FATE2. A. High proportion of genes with more than 5% unspliced reads (top) and near-uniform random distribution of unspliced proportions across genes (bottom) for datasets across mouse and human tissues. B. Projected velocity dynamics, indicating the differences between neighbourhood-based and direct projections; neighbourhood-based projection in UMAP, left; neighborhood-based projection in PCA, middle; directly-projected velocities in PCA, right. C. Schematic of metabolic labelling protocol and simplified cell treatment protocol in sci-FATE2 compared to sci-FATE, D. Low dimensional visualisations, providing broad view of key cell types and timepoints. Left; minimum distortion embedding of top 5 principal components, right; three-dimensional PCA (colours corresponding to cell types).E. Comparisons of sci-FATE and sci-FATE2; UMIs and genes per cell (left) and cell retention in chemical conversion protocol (right). F. Comparison of labelling and splicing data; across all genes (left) and for key genes in neural patterning (right).

This suggests that using a neighbourhood-based projection heuristic that alters velocity vectors by removing erroneous directions that cannot be reconstructed with the distribution of neighbouring cells potentially conceals inaccuracies in the high dimensional velocities inferred from splicing data.

### sciFATE2: An optimised metabolic labelling protocol

Observing the inconsistency of splicing data and issues with subsequent velocity inference, we considered metabolic labelling as an alternative, focussing on sci-FATE [20], as it offered the best reported combination of cellular throughput and technical requirements. To test this protocol, we performed a pilot (without 4sU labelling but with IAA treatment) which resulted in low data quality, with a median of approximately 1500 unique molecular identifiers (UMIs) per cell (Figure 1E, left).

To improve data quality, we made several changes to the protocol (Figure 1C), altering cell fixation and storage; RNAse inhibition; PCR amplification and library concentration and automating reverse transcription and PCR steps (along with barcoded primer addition). We also developed a simpler, less damaging IAA chemical conversion protocol that offers minimal impact on label detection with increases in UMIs per cell detected (Figure S1A) and reduction in cell loss (Figure 1E, right).

This optimised protocol provides a substantial improvement in data quality. With sci-FATE2, we capture around 14,000 UMIs per cell, representing nearly 5,000 genes per cell, from approximately 12,000 cells per experiment, and measured a doublet rate of 6% in a species-mixture experiment (Figure S1C). We observe specific detection of labelled signature in 15-20% of reads being detected as labelled (Figure 1F, left). By contrast, there was a nearly uniform random distribution of unspliced ratio across genes (Figure 1A, 1F), and 18.5% of genes lacked unspliced information, while only 0.2% of genes lacked labelling (Figure 1F).

With this optimised protocol, we set out to apply sci-FATE2 to examine the dynamics of patterning and differentiation in the developing neural tube. To this end, we used a protocol for directly differentiating mouse ES cells into neuromesodermal progenitors (NMPs), neural progenitors, and spinal cord neurons, using 500nM Shh agonist (SAG) to induce the most ventral neural tube identities (p3/V3, pMN/MN, and floor plate)[37, 38]. We labelled cells with 500*μ*M 4sU for 2 hours immediately prior to fixation (having confirmed that this dose led to no observable changes in transcriptional behaviour, Figure S2). Samples were taken on day 3 to 8 of the differentiation, during the period that NMPs and various neural cells are generated. All timepoints were prepared and sequenced together, with three full-time course replicates. A fourth replicate was collected, interpolating timepoints be-tween day 3 and 4 at five-hour intervals, to capture the rapid changes during NMP differentiation. After pre-processing, quality control and cell-type classification, we captured 44,713 time-resolved transcriptomic profiles that reflected the expected cell types and developmental progressions (Figure 1D).

### Velvet: deep generative velocity inference

To model gene expression dynamics, we focus on a generalised idea of ‘velocity’ that is not specific to splicing dynamics. We applied a biophysical model of RNA transcription, labelling and degradation as per previous studies [22, 25]. In this model, we can define a cell’s velocity in terms of three observables (labelled reads, total reads, labelling time) and one free parameter, *γ*, which represents a gene-specific degradation rate.

Even with this simple framework, inferring velocity and *γ* values across thousands of dimensions in gene expression space is a difficult task, and so to improve velocity inference, we developed a deep generative framework, velvet (Figure 2A). Motivated by the assumption that the dynamics exist on a lower-dimensional manifold within gene expression space,velvet learns a neural vector field embedded in the latent space of a variational autoencoder, mapping from the data to a ‘latent’ space (default 50 dimensions). This vector field is used to infer ‘latent velocities’, then total expression and velocity reconstructions (projected from the model’s latent space) are used to produce a reconstruction of the ‘new’ labelled expression. The loss function is defined to minimise the difference between the real and reconstructed total and labelled transcript datasets. The model thus learns i) a low dimensional representation of gene expression; ii) a vector field within this representation that captures cellular dynamics; iii) a biophysical equation that relates these ‘latent’ dynamics to the labelled-total relationships observed in the data.

**Figure 2:**
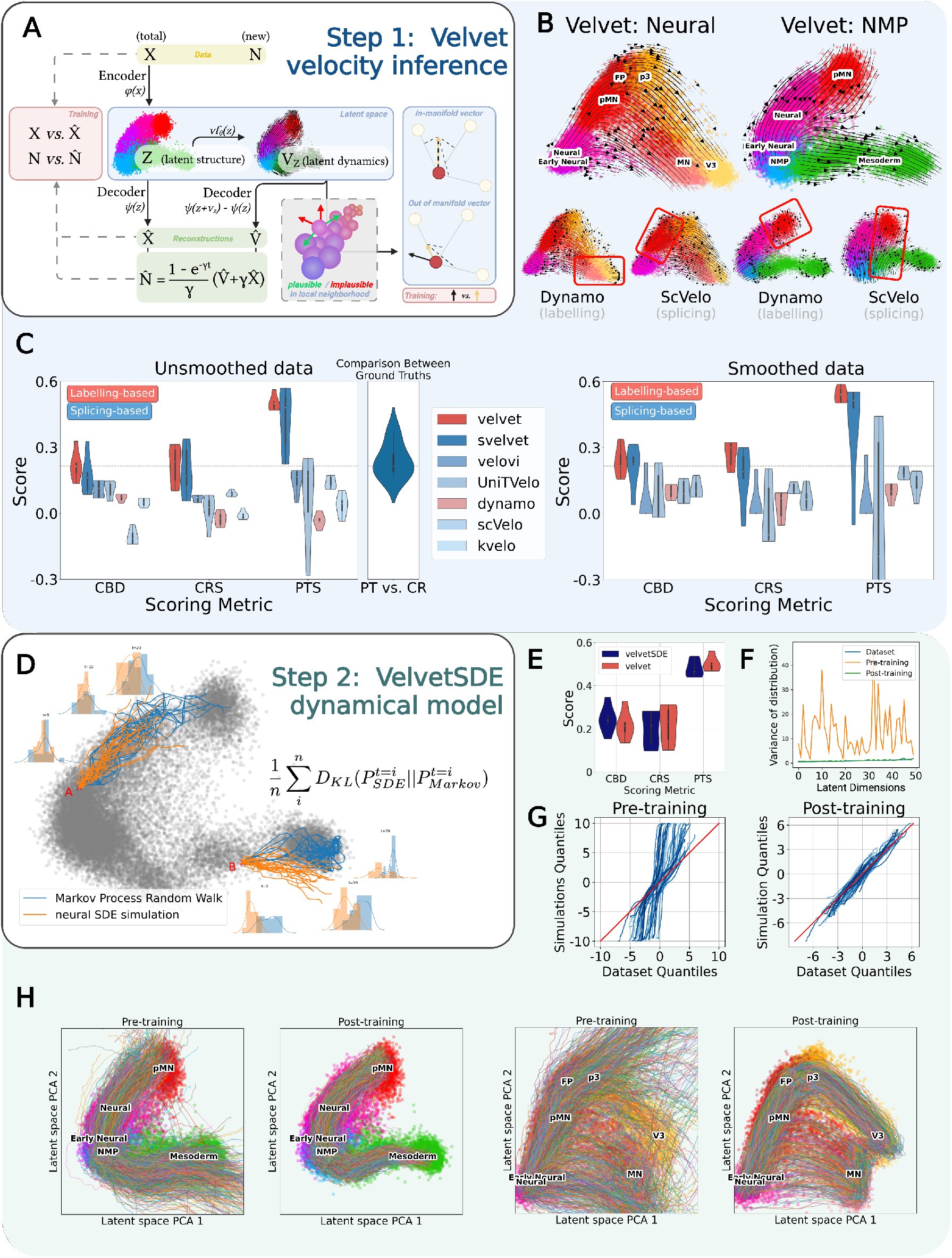
Dynamical modelling of expression dynamics with velvet & velvetSDE. A. Schematic of velvet module structure, demonstrating latent vector field, data reconstructions and neighbourhood constraint. B. PCA visualisation of velocity inferred from velvet, compared to dynamo (with labelling data) and scVelo (with splicing data). C. Quantitative benchmarking comparing velvet (and the splicing version, svelvet) against other velocity inference tools across six data subsets and three scoring metrics. D. Schematic of velvetSDE training: A demonstrates cell with good correspondence between nSDE and Markov simulations (orange and blue lines and histograms), B demonstrates cell with poor correspondence between nSDE and Markov, where training will update nSDE vector field. E. Benchmarking scores of trained velvetSDE compared to original velvet scores. F. Variance of pre- and post-training simulations across 50 latent dimensions, compared to variance of dataset. G. Q-Q plots comparing distributions of pre-trained simulations (left) and post-trained simulations (right) with the original dataset, compared in latent space. H. Comparison of velvetSDE stochastic trajectory simulation pre- and post-training.

To improve inference further, we reasoned that with sufficient sampling resolution, cells are expected to move towards other cells in their local neighbourhood — the exception being cells at a trajectory’s leading edges that are transitioning to unobserved states. In general, velocity analysis can be considered as a form of interpolation, describing the transition dynamics between observed states (as opposed to trying to describe transitions to unobserved states such as later cell types not captured in the dataset), meaning that when a velocity vector points away from all nearby cells in gene expression space, this is more likely to be an erroneous prediction than representative of the biology. As such, we add to velvet’s loss function a ‘neighbourhood constraint that penalises velocity predictions that point away from the convex hull of a cell’s nearest neighbours (details in methods). By constraining the directions in which each cell’s velocity vector can point, this penalty serves to reduce the solution space for the system’s dynamics, with the aim of improving the accuracy and robustness of velocity inference.

While neighbourhood information has previously been used in the projection of velocities for visualisation (Figure 1B), the neighbourhood constraint instead builds this information into the velocity inference process itself. Additionally, while it is common to incorporate neighbourhood information by performing ‘smoothing’ (replacing each cell’s expression with the mean expression of the local neighbourhood of cells), this constraint is designed to use the information in a more principled manner, explicitly based on the assumption of interpolative dynamics between observed states. Moreover, this neighbourhood constraint is a soft penalty that does not transform data as the above projection and smoothing methods do.

To assess whether velvet accurately captures known dynamics, we split the data into two observed decision systems: NMPs generating either neural or mesodermal cells; and neural progenitor cells differentiating into floor plate (FP), V3 interneurons and motor neurons (MNs). We inferred velocity for these decision systems separately. We found that velvet predicted dynamics that closely match the biology of the system, while other velocity tools (run with either labelling or splicing data) produce noisier predictions that less accurately described the known biology (despite using data smoothing, unlike velvet) (Figure 2B).

To assess performance quantitatively, we split the sci-FATE2 data into six subsets for benchmarking (ranging in size from 7000 to 45000 cells). First, we use cross-boundary direction correctness (CBD) [27], which assesses whether velocities correctly predict user-defined directionality between clusters. We additionally developed two parallel ground truths (Pseudotime score, PTS; CellRank score, CRS) based on the known dynamics of each data subset, using the trajectory analysis tools scFates and CellRank [41, 42]. These ground truths test the degree to which velvet can accurately describe the coarse cell state transitions known to occur in the data.

We compared velvet to five velocity tools [18, 25, 27, 28, 43] (four using splicing, one using labelling) and found velvet consistently outperformed all other approaches across the three metrics (Figure 2C). Moreover, velvet performed as well without any data smoothing, a step conventionally required of velocity analysis that can remove biological signal. We found velvet scored closely to a positive control comparing the similarity of PTS and CRS ground truths, suggesting that the performance of velvet is achieving maximal scores for the ground truth comparisons used, in contrast to all other velocity tool tested.

Extending the model to handle splicing data (with ‘svelvet’, details in Methods), we find that our approach still outperforms other methods across all metrics with splicing data. This suggests that svelvet could be of use for existing RNA velocity datasets - however, we note that this approach is not designed to handle or overcome confounding factors that can be present in the splicing dynamics of different datasets [17, 44, 45]. With splicing data, we saw lower and more variable scores with splicing data than with labelling data (Figure 2C), consistent with the idea that metabolic labelling provides more consistent, higher quality temporal information.

### VelvetSDE: dynamical modelling with neural SDEs

With velvet’s improved velocity inference, we reasoned we may be able to use inferred dynamics to simulate cellular trajectories for dynamical analysis of differentiation. However, we found that trajectories simulated from a system’s initial state failed to reach expected terminal states, instead being predicted to move to areas where no data points are observed (Figure 2I).

We hypothesised that the problem was that the inference process considered no long-term information. Nothing in the framework of instantaneous velocity estimation encodes the requirement that cells should maintain their trajectories within the data manifold, or that initial states should progress to reach terminal states. As such, errors that are negligible within the context of a single timestep may accumulate across successive timesteps to produce large deviations. To solve this, the global dynamics of the system must be modelled, extending pointwise velocity predictions to a system of differential equations that describe how cell states evolve over longer timescales.

From this perspective, a developmental system can be considered a random process described by drift-diffusion stochastic differential equations (SDEs). A common approach to approximate such a drift-diffusion model with single-cell data is to draw upon the equivalency of drift-diffusion models and random walk operators on a graph [7, 42, 46]. Through this approach, stochastic trajectories can be simulated as random walks across the cell-cell similarity graph, with transition probabilities calculated based on inferred velocity dynamics. However, this method has drawbacks: stochasticity arises from random jumps between cells/states, providing a simplistic noise model that cannot be refined or adjusted. The resulting trajectories are noisy and discontinuous (Figure S6A).

Instead, we used the recently developed neural stochastic differential equation (nSDE) framework [35, 36] to model a drift-diffusion system. In this framework, the SDE is a neural network with ‘black-box’ parameters, and numerical integration of the system is compatible with back-propagation and gradient descent. In velvetSDE, the drift component is the velvet vector field, and diffusion is represented by a random walk of equal magnitude in all directions (i.e. scalar diagonal Brownian motion).

To train the nSDE, we take advantage of the drift-diffusion equivalency with Markov random walks with the following steps (Figure 2D): i) set the noise magnitude of the nSDE system to approximate the noise observed in Markov walk simulations, ii) generate nSDE distributions and Markov random walk distributions for each cell, iii) define the model’s loss function as the Kullback-Leibler divergence of nSDE trajectory distributions from Markov trajectory distributions (assuming gaussian distribution across a timestep of a simulation’s trajectories).

This training step modifies the vector field to produce on-data trajectories. However, since both Markov and nSDE simulations are based on the same inferred velocities, this training is not expected to significantly alter the overall dynamics of the system. The stochasticity of the nSDE framework facilitates comparisons with Markov random walks - but after training, the nSDE’s noise function can be adjusted freely, allowing the effects of noise to be assessed in a manner not possible with simulations based on Markov random walks.

VelvetSDE trajectories show significant improvements in staying within the data distribution (Figure 2I) without altering benchmarking scores (Figure 2E), suggesting that underlying biological dynamics are conserved. Pre-training simulations show a high degree of variance across latent dimensions, and poor concordance with the latent distribution of the dataset; but after velvetSDE training, simulations have variance that closely matches that of the dataset (Figure 2F), and distributions show strong concordance between data and simulation (Figure 2G, Figure S3E). These results suggests that, through the nSDE framework, we can learn a dynamical model that can be used to simulate trajectories that faithfully recreate the distributions of the underlying data.

### Predicting cell fate dynamics in NMPs

To explore velvetSDE’s dynamical predictions, we first focused on cell fate decisions in NMPs. Simulating all D3.2 (Day 3 + 5 hours) cells (with a noise magnitude of 0.1, used as default unless otherwise specified), we found resultant trajectories clustered into neural and mesodermal fates (Figure 3A). We reasoned that with a single-cell mapping of cell fates, we could define the ‘decision boundary’ between fate-committed regions, which could be used to analyse temporal and genetic factors relating to cells’ potency through differentiation. To examine the boundary between fates, we found cells that had varying fate predictions over multiple simulations: we generated stochastic fate distributions by simulating each cell’s trajectory 10 times (chosen arbitrarily; changing this number affects the can affect the size of mixed-fate population, as can changing the model’s noise magnitude). Each cell’s distribution could then be classified as entirely ‘neural’, entirely ‘mesoderm’, or a ‘mixed’ distribution that contained some neural and some mesodermal trajectories (Figure 3B). PCA visualisation of these classifications in velvetSDE’s latent space revealed distinct decision boundaries separating ‘mixed’ progenitors from lineage restricted progenitors (Figure 3C). We found this boundary, which reflects the commitment point between these two fates, aligned with the boundary between expression and velocity dynamics of known markers of neural, NMP and mesodermal cell types (Figure 3E), indicating that this mixed-fate population reflects an intermediate region in gene expression space across the variations in cell states and expression dynamics in the D3.2 population.

**Figure 3:**
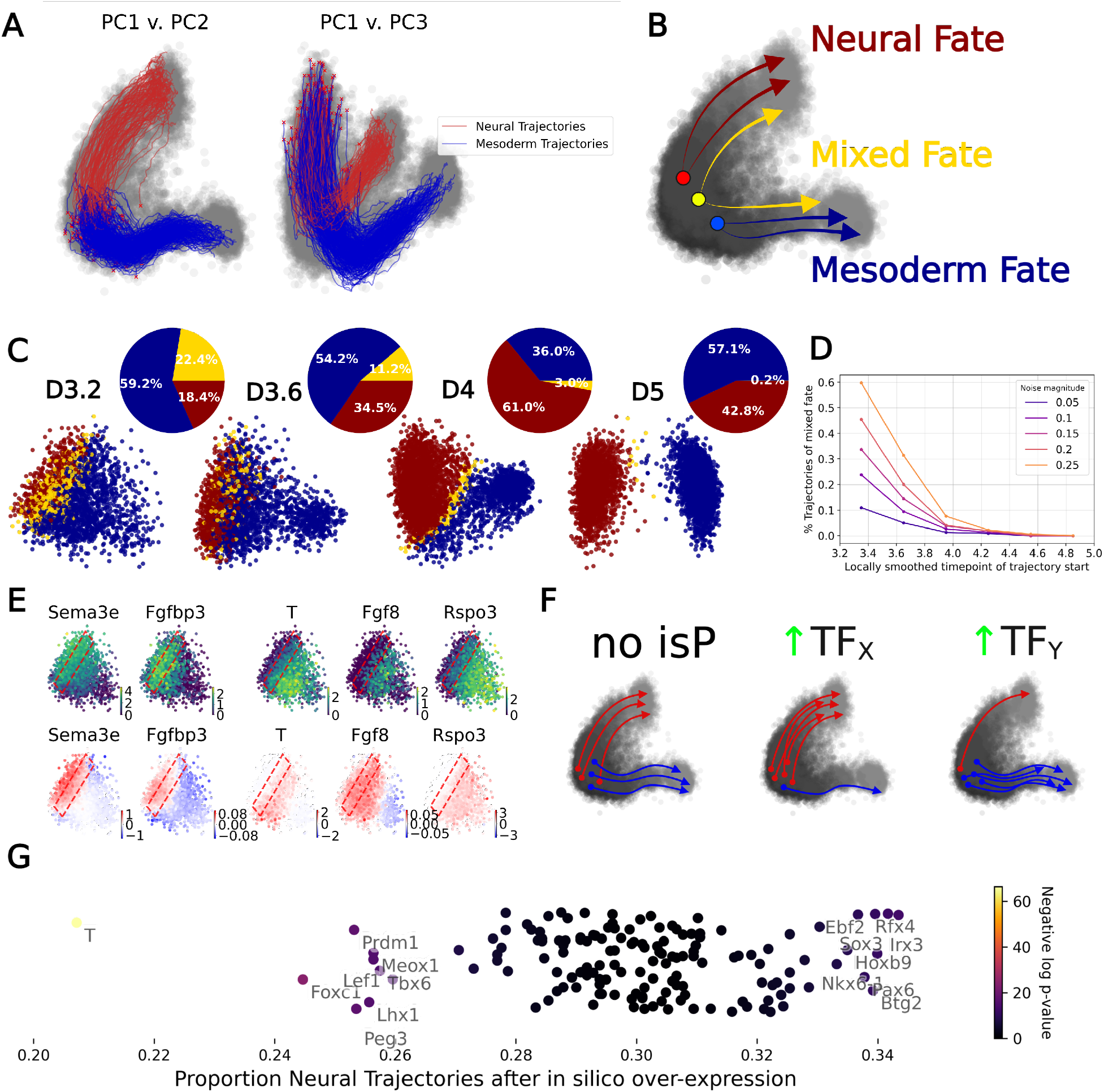
Modelling the decision boundary between fates in NMPs. A. Trajectories from D3.2 NMPs, clustered into two distinct fates: neural and mesodermal. B. Schematic demonstrating the concept of stochastic fate simulation and mixed-fate cells. C. Latent space PCA of cells across timepoints, coloured by their predicted fate; entirely neural (red), entirely mesodermal (blue) or a mixture of the two (yellow; decision boundary), with pie charts representing the proportions of each group. D. Proportion of cells predicted to have a ‘mixed’ fate, as a function of time window and noise magnitude of velvetSDE model. E. Expression and predicted velocity of neural, NMP and mesodermal markers, red box highlighting the decision boundary region. F. Schematic for in silico perturbation (isP) test for key genes in the axes of the decision boundary G. results of isP test showing top pro- and anti-neural hits visualised.

We examined how the mixed population evolved over time. The proportion of mixed fates decreased monotonically until none were left by day five (Figure 3C). To examine the effect of velvetSDE’s noise on this mixed-fate population, we repeated trajectory simulation across a range of noise magnitudes and observed that the mixed cells had largely disappeared by day 4 (Figure 3D), consistent with the observation that neural commitment TFs such as Olig2 are induced on day 4 [38].

To test further the validity of the inferred decision boundaries, we identified the transcription factors (TFs) that had the largest effect on the boundary. For each TF in the data, we set expression to the maximum observed value across all mixed-fate cells, re-projected these perturbed data points to the velvetSDE latent space and simulated each perturbed cell’s trajectory to observe how in silico perturbation affects the predicted distributions of fates (Figure 3F). Ranking genes by the magnitude of their effect, we see the highest ranked hits for increasing or decreasing the ‘neural’ proportion of fates include many known TFs known to play a role in neural or mesodermal specification (Figure 3G). Visualising the loading contribution of top hits to the model’s latent space, we see these genes broadly correspond to the observed dimension of the neural-mesodermal decision (Figure S4C).

In summary, stochastic simulation of trajectory distributions delineates fate commitment points and identifies fate specifying factors. This approach provides a highly resolved picture of cell fate decision structure and timing, and facilitates the generation of hypotheses regarding fate commitment, potency time-windows, and potential reprogramming factors that can be subsequently tested through genetic perturbation and lineage tracing.

### Examining gene expression dynamics in neural cells

We next examined the trajectory distributions generated by velvetSDE, concentrating on neural cells. Clustering simulations of day 4 early neural cells revealed three distinct fates: motor neuron (MN), V3 interneuron (V3), and floor plate (FP) (Figure 4A). Predicted trajectories can be projected to gene expression space to produce distributions of time series that exhibit the expected median dynamics, but with considerably variation between individual trajectories (Figure 4B).

**Figure 4:**
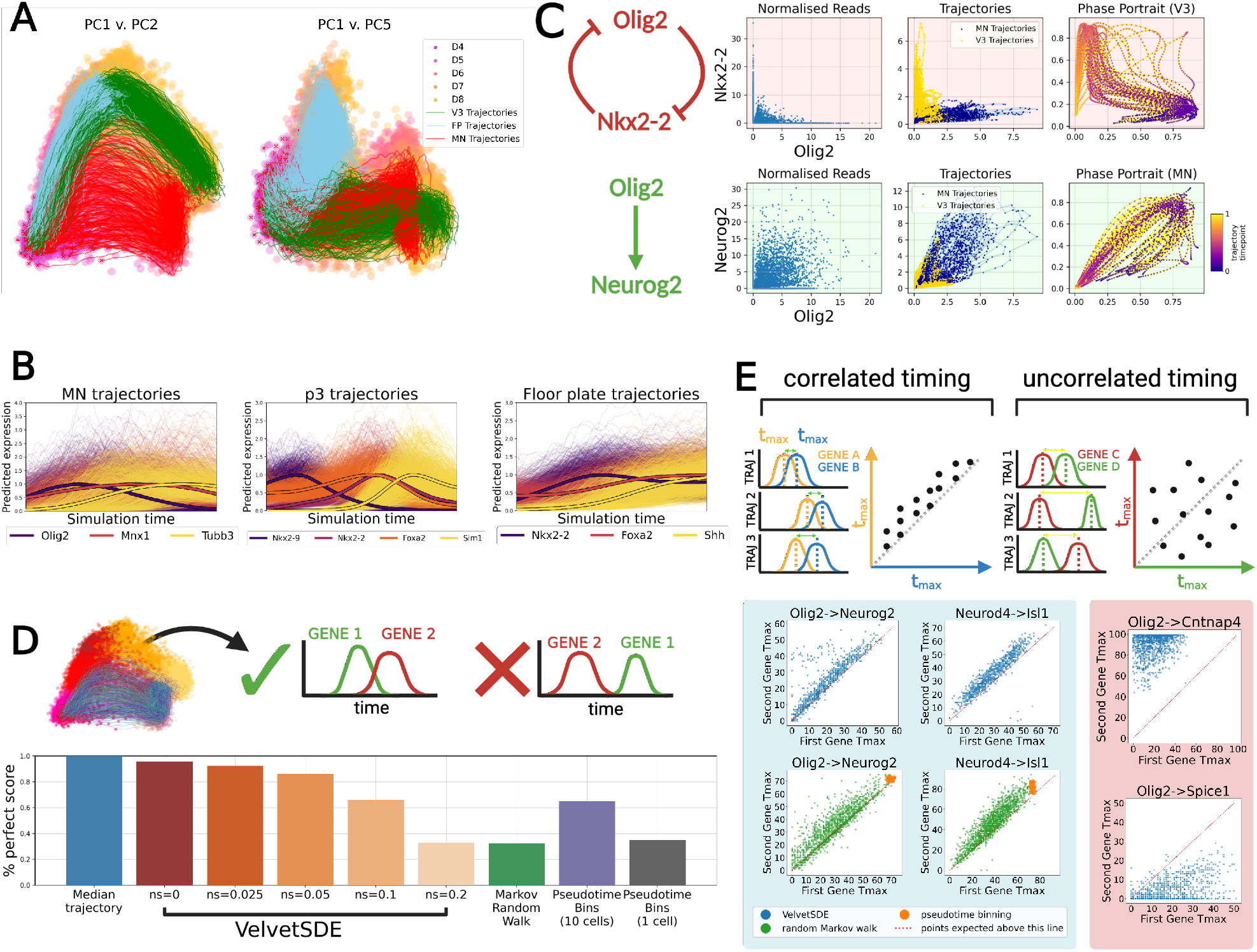
Dynamics and structure of gene expression in neural cells. A. Simulation of early neural cell trajectories, clustered into motor neuron (MN), V3 interneuron (V3) and floor plate (FP) fates. B. Gene expression averages and distributions, mapped from simulated trajectories, for the three fate clusters. C. Comparison of normalised reads data and phase portraits of normalised, gaussian trajectory distributions for example interactions: cross-repressive (Olig2 & Nkx2-2) and directly causal (Olig2 & Neurog2). D. Scoring trajectory distributions for the conservation of known biology, defined as a series of seven known gene orderings (e.g. Olig2 before Neurog2) and measured by comparing the time of maximal expression across trajectories; score represents the percentage of trajectories that perfectly capture all orderings. E. Correlation of time dynamics analysis: top shows schematic illustrating the concept. VelvetSDE simulations for related genes in blue box, top. Markov random walk and pseudotime binning controls shown in blue box, bottom (green and orange respectively). VelvetSDE for unrelated genes shown in red box.

To explore the dynamical information contained in these trajectories, we focussed on two example interactions, a direct cross-repressive interaction (Olig2 and Nkx2-2)[47] and a directly causal activation (Olig2 and Neurog2)[38, 48]. With normalised reads alone, we see a noisy picture: the difference between these two relationships is discernible only through the degree to which the the gene pairs are co-expressed (Figure 4C, left). With predicted trajectories, we can provide more detail, visualising the different dynamics observed in different fate trajectories (Figure 4C, middle). Visualising phase portrait plots (through normalisation and Gaussian smoothing of trajectories), we can get further insight into the different relationships, visualising the consistent structure of sequential co-expression of Olig2 and Neurog2 and mutual exclusivity of Olig2 and Nkx2-2 (Figure 4C).

We next looked to test whether biological information is conserved across the variability of predicted gene expression time series distributions. Focusing on motor neuron trajectories, we defined a set of expected expression orderings (for example, maximum Olig2 expression should precede maximum Neurog2 expression [38]). We then calculated the percentage of simulated trajectories that capture all orderings correctly. We compared velvetSDE simulations with different noise magnitudes to three baseline models representing conventional tools of single-cell: i) a Markov random walk model with transition matrix based on velocity dynamics, ii) binning cells by pseudotime values (also based on velocity dynamics and calculated with scVelo [43] and choosing 10 cells per bin, and iii) pseudotime binning with 1 cell per bin. With no noise, velvetSDE simulations almost perfectly conserve expected gene orderings (Figure 4D).

As the noise magnitude increases, scores decreases gradually to reach a level equivalent to Markov random walks. Pseudotime binning performed better when averaging ten cells per bin, but selecting individual cells scored poorly.

However, a unique potential of velvetSDE simulations is not only to preserve biological information, but also to capture meaningful variation across the different simulated trajectories of a fate that would be lost in averaged or pseudo-temporal trajectories. To explore this, we continued to use a gene’s time of maximal expression, *t*_*max*_, as a metric to explore variation across trajectories. Focussing on VelvetSDE trajectories with 0.1 magnitude noise, plotting the distribution of *t*_*max*_ values for pairs of genes, we saw a high degree of variation in the time of maximal expression for different genes. Importantly, these distributions are highly correlated for related gene pairs (genes known to be involved in the temporal progression of motor neuron differentiation, such as Olig2 and Neurog2 [2, 38]) (Figure 4E, blue box, top) while for unrelated genes (Olig2 and randomly chosen genes in the data, Cntnap4 and Spice1), they are not (Figure 4E, red box).

We see that Markov random walks capture similar covariation - however with slightly more variability leading to more incorrect orderings (Figure 4E, green points; incorrect orderings appear below red dotted line). With pseudotime binning, the observed covariation in *t*_*max*_ is entirely lost (Figure 4E, orange points overlain on green points), which may result from the loss of variability through averaging per bin (plotting pseudotime-binned trajectories supports this idea, Figure S6A). As a control, we compared velvetSDE simulations to trajectories from a ‘technical noise’ model where all variation is from from differences in initial conditions and the addition of random noise, finding that the technical noise model failed to capture the correlative structure in expression timings, indicating that the correlation is biologically meaningful (Figure S6B).

VelvetSDE trajectory distributions appear to simultaneously preserve core biological information while capturing variation between trajectories and correlation between related genes. This correlative structure is lost in averaging and cannot be explained by technical noise, suggesting that velvetSDE’s modelling framework offers a valuable tool for studying gene regulatory dynamics.

### Applying velvet to the dynamics of in vitro patterning

Finally, we applied our data and framework to the question of neural fate decisions in vitro. In our experiments we observed the formation of three ventral neural tube fates — floor plate (FP), V3 interneuron (V3) and motor neuron (MN). In vivo, these fates are exposed to different concentration and duration of Sonic Hedgehog (Shh) signalling [49]. However, in vitro, all cells are exposed to the same concentration and timing of Shh agonist treatment (500nM daily from day 3). This raises the question of what dynamics drive the formation of these distinct fates within homogeneous condition; whether they form stochastically through varying interpretation of signalling cues or appear sequentially due to prolonged exposure to Shh signalling. We reasoned that with the increased dynamical resolution of our data and modelling framework, we might be able to provide insight into the dynamics of this patterning decision.

We took trajectory distributions simulated from day 4 early neural progenitors and clustered into MN, V3 and FP fates (Figure 4A). Observing that motor neurons arise before V3 interneurons (consistent with previous studies [50]), we first asked whether Olig2 expression is as high in progenitors that become V3 as those that become MN, i.e. whether p3 progenitors arise sequentially from Olig2+ pMN progenitors, or in parallel but at a later timepoint. Plotting the expression of marker genes (Irx3, early neural; Olig2, pMN; Nkx2-2 p3; Foxa2, floor plate; Sox2, pan-neural progenitor), indicated that Olig2 expression was detectable in V3 and FP trajectories, but at a lower level than in MN trajectories, suggesting that high levels of Olig2 expression is associated commitment to MN fate (Figure 5A), consistent with previous biological observations [38].

**Figure 5:**
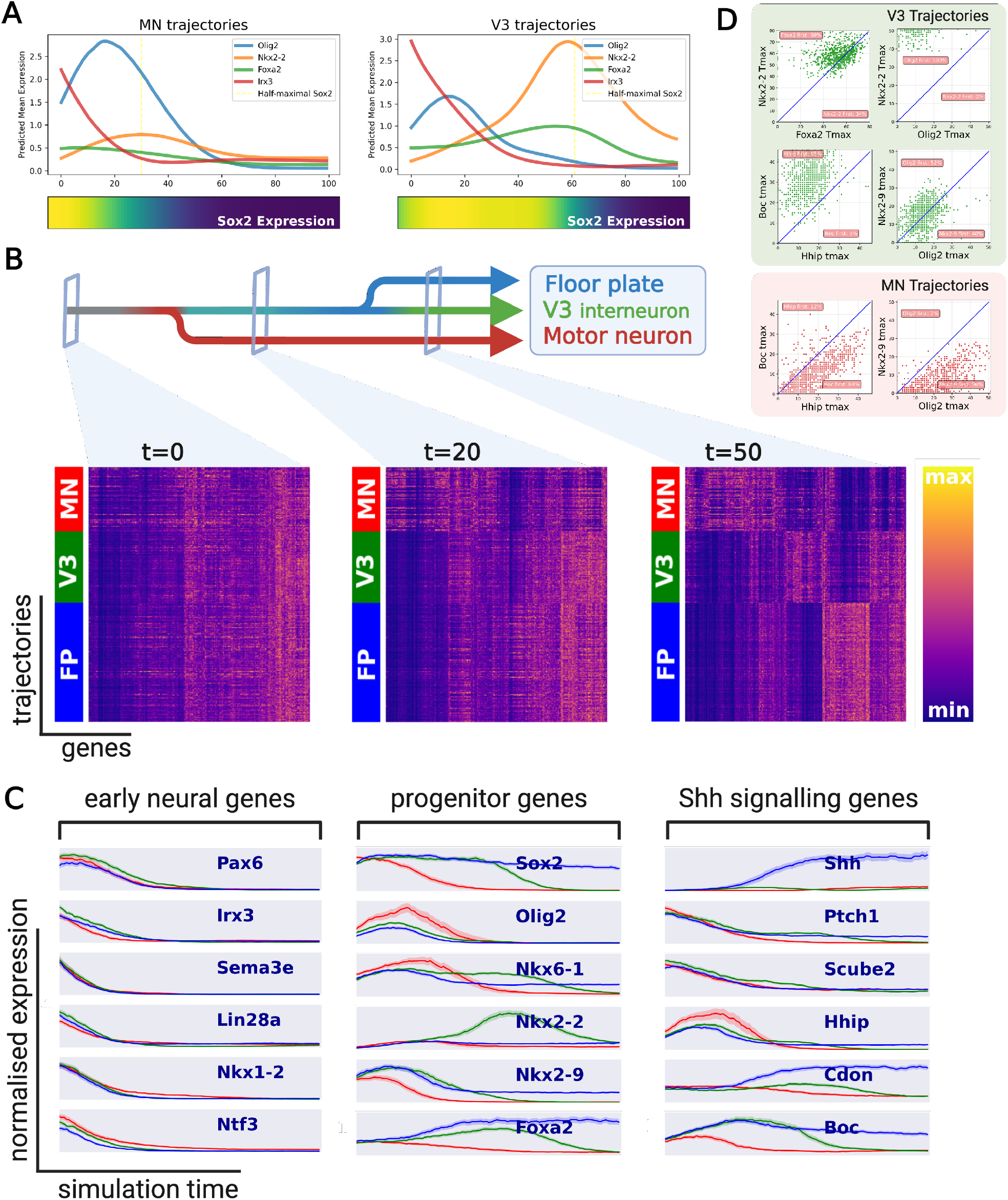
Decision dynamics in neural patterning in vitro. A. Average expression profiles for key genes in MN trajectories (left) and V3 trajectories (right), B. Clustermaps of all genes across trajectories across three fates at specified timepoints. Genes are clustered; trajectories are maintained in their fate clusters. C. Median trajectory from 100 randomly sampled trajectories for early neural genes (left), progenitor markers (middle) and Shh signalling genes (right), showing expression value for three trajectories: MN (red), V3 (green), FP (blue). Shaded area represents the 99% confidence interval across trajectories. D. Expression timing analysis for key genes in V3 trajectories (green) and motor neuron trajectories (red).

Analysing the timing of patterning decisions through conventional single-cell analysis can be challenging because cells with different fates are difficult to distinguish and a range of different developmental times can exist within a single collected time-point. With VelvetSDE simulations, we can cluster early timepoints based on their predicted downstream fates, and we can view cross-sections of these trajectories at a particular timepoint, allowing the dynamics of differentiation to be analysed in more detail. To explore the timing of neural patterning decisions, we produced clustermaps to visualise all genes of all trajectories at particular timepoint cross-sections, grouping by fate. Initially, there is very little to distinguish the three fates (Figure 5B, left). By an early stage (20 timesteps of 100), the gene expression profile of MN trajectories was distinguishable from V3 and FP trajectories in the clustermap, but these latter fates were less distinct (Figure 5B, middle). Subsequently, the three fates could be clearly distinguished (Figure 5B, right).

Examining median trajectories plots of key genes for the three fates indicated that many early neural markers, such as Pax6 and Irx3, have highly consistent expression profiles across the fates (Figure 5C, early genes), suggesting that the exit from an ‘early neural’ state is similar across the three fates. Contrary to this synchronised pattern of early neural genes, subsequent neural progenitor markers show clear distinctions between fates, starting with Olig2 and Nkx2-9 distinguishing MN from non-MN fates (Figure 5C, progenitor genes), consistent with the role of Nkx2-9 in the specification of both FP and V3 cell types [51, 52]. Of note, while Olig2 and Nkx2-9 appear to be the earliest key marker genes to separate MN from the other fates, the earliest distinction between V3 and FP fates appears to be the induction of Shh itself in FP (Figure 5C, Shh genes).

Additionally, we observed variation in the expression of Shh signalling regulators between the fates: Hhip, a negative regulator of Shh [53], expression appears higher in MN trajectories and Boc, a Shh coreceptor [53], expression is higher in FP and V3 fates (Figure 5c, Shh genes).

A comparison of the time of maximal expression for marker genes shows Olig2 is always expressed before Nkx2-2 in V3 trajectories, but less consistently so for Nkx2-9, while peak Foxa2 expression is roughly concurrent with Nkx2-2, consistent with the above observations (Figure 5D). Additionally, we see the expression patterns for Olig2-Nkx2-9 and Hhip-Boc differ considerably in MN trajectories and V3 trajectories.

The dynamics described by this model are consistent with a two-stage decision: first MN vs. FP/V3, followed by a V3 vs. FP decision. This is consistent with previous findings regarding the timing of differentiations and expression patterns of key genes, and also corresponds well with recent findings that chromatin landscape changes specifically distinguish FP and V3 cells from more dorsal neural tube domains [54].

The time delay between MN fates and V3/FP fates could correspond to exposure to additional doses of Shh agonist prompting cells that are still Shh-responsive (i.e. cells that have not yet expressed sufficient Olig2 to differentiate into motor neurons) to shift from heading towards a MN state to a V3/FP state. As such, future work could explore whether changes in Shh signalling at critical timepoints (for example, around D5D6) could specifically bias differentiations either towards or away from a MN fate.

However, neural fate decisions do not appear to be entirely sequential or solely down to Olig2 levels, with Nkx2-9 and Shh regulators Boc and Hhip also showing early differences in expression between fates. While this observation is consistent with previous findings recording the complex differential regulation of Shh co-receptors such as Boc [55], it is not clear whether variation in Shh regulators is a cause or consequence of Olig2 and Nkx2-9 dynamics. These findings suggest that variability in signal interpretation may be involved in the different fate decisions observed in vitro, and that specific targeting of Shh regulators could provide an approach to further control the course of fate decisions in vitro.

## DISCUSSION

Here we have developed an integrated framework of experimental and computational methods for dynamical modelling of gene expression from time-resolved transcriptomics data. While splicing and metabolic labelling data are both regularly used for the inference of single-cell temporal dynamics, we found that the signal quality of splicing data is insufficient, while the data quality of a recently developed metabolic labelling protocol was low. Our optimised, semi-automated protocol, sci-FATE2, provides a controllable, consistent temporal signal with data throughput and quality comparable to commercial platforms. To infer dynamics with sci-FATE2 data, we developed a velocity inference method, velvet, which builds local neighbourhood information into the inference process of a mechanistic, generative model, resulting in velocity predictions that appear more accurate and score consistently higher in benchmarking than existing methods. To reconstruct overall system dynamics, we extended velvet using neural stochastic differential equation approaches, exploiting qualities of Markovian random walks to constrain trajectories to the data. With this, we were able to simulate trajectories of initial cells through to terminal states to produce simulations that accurately recreate the distributions of the underlying data and match known biology.

This extended approach, velvetSDE, allows new forms of analysis: simulating stochastic fate distributions across cells allowed the prediction of decision boundaries between fates, while trajectory distributions capture potentially informative biological variation in the temporal dynamics of gene expression. Applying the approach to the question of a cell fate specification in vitro, we were able to apply velvetSDE’s dynamical perspective to describe the dynamics of sequential fate decision making and resolve fate-specific variation in Shh regulation.

The throughput of sci-FATE2 can be increased with additional rounds of indexing [56], while further extensions of labelling (e.g., additional thio-nucleotides[57], varying labelling durations), alongside optimisations of the chemical conversion protocol and computational methods for label detection could offer further improvements to data quality and the temporal signal.

While velvetSDE is honed by training trajectories alongside Markov random walks, we found that the neural SDE framework allowed a greater control of stochasticity: without noise, velvetSDE simulations accurately preserved known biology, while the introduction of noise allows the analysis of boundaries between fate-restricted regions of progenitor cells. Importantly, velvetSDE’s framework opens up the possibility for future work to learn more sophisticated and biologically-motivated models of noise and uncertainty, allowing analyses that robustly handle stochasticity as a meaningful part of the biological systems we study.

We expect the framework of metabolic labelling and dynamical modelling to provide greater resolution for examining the dynamical effect of perturbations in developing systems. It also offers a natural connection between velocity-based analysis and lineage tracing methods. Future implementations could seek to extend this approach into a unified model that handles labelling or splicing information alongside longer-term lineage information.

Through the integration of temporal transcriptomics and mechanistic generative modelling, we can capture high quality information on the dynamics of expression, allowing new forms of analysis into cell fate decisions and providing a dynamical framework for modelling complex regulatory mechanisms in development.

## ACKNOWLEDGEMENTS

We are grateful to Fabian Fröhlich, Jake Cornwall-Scoones, Alberto Pezzotta and David Rand for fruitful discussions, as well as Ashley Libby, Tiago Rito and other members of the Briscoe lab for their constructive comments on the manuscript. We are grateful to the Advanced Sequencing STP and Flow Cytometry STP at the Francis Crick Institute for their assistance in carrying out experiments. This work was supported by the Francis Crick Institute which receives its core funding from Cancer Research UK (CC001051), the UK Medical Research Council (CC001051), and the Wellcome Trust (CC001051); by the European Research Council under European Union (EU) Horizon 2020 research and innovation program grant 742138 and by the Wellcome Trust (220379/D/20/Z). For the purpose of Open Access, the author has applied a CC BY public copyright licence to any Author Accepted Manuscript version arising from this submission.

## AUTHOR CONTRIBUTIONS

R.J.M and J.B conceived the project, interpreted data and wrote the manuscript. R.J.M. designed and performed experiments, developed the modelling framework, performed computational and data analysis.

R.J.M. and D.S. developed and optimised the automation of the experimental protocol.

## METHODS

### Cell culture and differentiation

Cells were cultured and differentiated as described previously [37, 38, 54]. HM1 (Thermo scientific) ES cells were maintained in ES cell medium with 1,000U/ml LIF on mouse embryonic feeder cells (mitotically inactivated). These cells were dissociated in 0.05% Trypsin (Gibco) and plated on tissue culture plates 20 minutes two successive times to remove feeder cells. Remaining supernatant cells were then plated on CellBind six-well plates (Corning) pre-coated with 0.1% gelatine solution, 60,000 cells per well in 1.5ml N2B27 + 10ng/ml bFGF. On day 2, the medium was replaced with N2B27 with 10ng/ml bFGF and 5*μ*M CHIR99021 (Axon) for 20 hours. Subsequently and every 24 hours afterwards, medium was replaced with N2B27 with 100nM RA (Sigma) and 500nM SAG (Calbiochem).

## Cell labelling, collection, fixation and storage

Cells were incubated in 500*μ*M 4sU (Sigma, made into 500mM stock dissolved in DMSO) for two hours in the appropriate medium in the dark. Collection was then done as swiftly as possible to minimise the time between labelling and fixation. For this, cells were washed in PBS, dissociated in 500*μ*l room temperature accutase for two minutes, spun at 1000g for 3 minutes. For fixation, cells were resuspended in 400*μ*l PBS + 0.1% DEPC (Sigma) + 10mM DTT (Sigma) and 1600*μ*l Methanol + 0.1% DEPC + 10mM DTT was added slowly, drop-wise to cells. Cells were kept on ice, rocking at 20rpm for 30 minutes, and were then stored at -80^*°*^C for up to three weeks. Note that the melting point of methanol is -97.6^*°*^C, and as such this fixation protocol will not result in freeze- or thaw-associated damage to cells. The number of wells used for a single fixed sample was chosen such that 5-10 million cells were fixed per sample.

## sci-FATE2

Summary of main protocol changes: Noting that PFA fixation, Triton-X and freeze-thaw cycles are all associated with increased RNA damage or loss [58, 59], we adopt methanol fixation and liquid-phase storage at - 80°C. Finding that enzymatic RNAse inhibitors are ineffective [60], we add DEPC to our fixation and wash buffers. To eliminate manual errors and reduce time spent with samples on ice, we automated reverse transcription and PCR steps with the Mosquito HV (SPT Labtech). To minimise over-amplification, we reduced the number of PCR cycles from twenty to ten, and concentrated the library with SPRI beads. To reduce the harshness of the IAA chemical conversion step, we removed the use of hydrochloric acid and 50% DMSO, finding that doing so did not negatively impact the detection of labelled reads or the total number of UMIs per cell (FIGURE), and provided an increase in cellular retention through the protocol (figure 1H)

For chemical conversion, cells were kept on ice at all times and all spins were done at 2000rpm, 4°C for 5 minutes. Cells were first moved from -80°C onto ice for 3 minutes and gently resuspended. Cells were then spun down and resuspended in 1ml PBS + 0.1% DEPC + 3% v/v dissolved BSA solution (NEB) + 10mM DTT, spun again and resuspended in 100*μ*l PBS + 3% v/v dissolved BSA solution. To this suspension, 220*μ*l water, then 40*μ*l sodium phosphate buffer then 40*μ*l 100mM IAA (Sigma, dissolved in ethanol) were added. Cells were incubated at 50°C for 15 minutes, being gently resuspended every five minutes. Cells were then added to a quenching mix of 1.5ml PBS + 3% v/v dissolved BSA solution and 5mM DTT to quench the chemical conversion reactions. Cells were spun down and resuspended in PBS at a concentration of 1 million cells per ml. Sci-FATE2 library preparation was performed immediately. Cells were distributed 2*μ*l per well into a LoBind 384-well plate using Mosquito HV (SPT Labtech). 1*μ*l of oligo-dT primer (5’-ACGACGCTCTTCCGATCTNNNNNNNN[10-bp well-specific barcode]TTTTTTTTTTTTTTTTTTTT-TTTTTTTTTTVN’-3, Integrated DNA Technologies) was added to each well with Mosquito, and plate was heated at 55°C for five minutes before being immediately placed on ice for two minutes. To each well, 2*μ*l of first-strand reaction mix (1*μ*l SSIV buffer, 0.125*μ*l DTT 10mM, 0.125*μ*l dNTPs 10mM, 0.125*μ*l RNase inhibitor and 0.125*μ*l SSIV reverse transcriptase) was added to each well with Mosquito, and the plate was put through the following reverse transcription thermal schedule: 4°C for 2min, 10°C for 2min, 20°C for 2min, 30°C for 2min, 40°C for 2min, 50°C for 2min and 55°C for 10min. Cells were then pooled manually, mixing well to ensure cells are resuspended. To cells, 1.8*μ*l of 1mg/ml DAPI (ThermoFisher) was added. 50 cells per well were then sorted into four 96-well plates with 3.6*μ*l water and 0.4 second strand synthesis buffer in each well. This was done with a MoFlo XDP, with a 100*μ*m nozzle, gating on scatter and DAPI to stringently remove doublets and debris. To each well of each plate, 1*μ*l of second strand synthesis mix (0.65*μ*l water + 0.1*μ*l second strand synthesis buffer + 0.25*μ*l second strand synthesis enzyme) was added, plate were vortexed at approximately 1000rpm for five seconds and spun down for five seconds, and were then incubated at 16°C for three hours. Plates were then stored with foil lids at -80°C for up to a month. Tagmentation and clean up were performed manually, two plates at a time. For tagmentation, plates were thawed and 5*μ*l tagmentation mix (4.875*μ*l tagmentation buffer + 0.125*μ*l Tn5 4nM) was added. Plates were incubated at 55°C for five minutes, before being incubated at room temperature for five minutes with 10*μ*l DNA-binding buffer (Zymo) to quench the reaction and lyse cells. For clean up, 30*μ*l Ampure SPRI beads (Beckmann Coulter) was added to each well. Plates were vortexed at 1000rpm for 10 seconds and incubated at room temperature for five minutes. Plates were added to magnets, 42*μ*l supernatant was removed and each well was washed with 100*μ*l 80% ethanol twice. Plates removed from magnets and left to air dry for 2-3 minutes. 10*μ*l EB buffer was added to each well, and plates were vortexed at 2000rpm for one minute and spun briefly for seven seconds, before being left to incubate at room temperature for ten minutes. Plates were returned to magnets, and 9*μ*l su-pernatant was moved to a 384-well Lobind plate. 1*μ*l PCR primer mix containing 10*μ*M P5 primer (5’-AAT GATACGGCGACCACCGAGATCTACAC[i5]ACACT-CTTTCCCTACACGACGCTCTTCCGATCT-3’; IDT) and 10*μ*M P7 primer (5’-CAAGCAGAAGACGGC-ATACGAGAT[i7]GTCTCGTGGGCTCGG-3’; IDT) was added to each well of 384-well plate with the Mosquito, and 10*μ*l NEBNext PCR mix was added manually. PCR was performed with the following program: 2C for 5min, 98C for 30s, and 10 cycles of 98°C for 10s, 66°C for 30s, 72°C for 1min) and a final 72°C for 5min. The full library was then pooled for a total volume of around 7ml. This library was distributed into 8 samples of 850*μ*l. For each, an Ampure SPRI clean up was performed with 0.9X beads, eluting each sample in 100*μ*l. These eluted samples were collected into two 400*μ*l samples. To these samples, double size selection was then performed: 200*μ*l SPRI beads were added, incubated for ten minutes and then added to a magnet. 600*μ*l of supernatant was moved to new tubes, where 160*μ*l fresh beads were added to sample. After 5 minutes of incubation, samples were again added to magnet, and washed twice with 1ml 80% ethanol. The two samples were each eluted in 30*μ*l for 60*μ*l total library product. The final library product was quantified using Qubit and BioAnalyzer. For pilots, libraries were sequenced on NovaSeq SP with 700m reads per sample, configuration: 130-10-10-18. For experiments, libraries were sequenced two samples at a time on NovaSeq S1, total approximately 900m reads per sample (75,000 raw reads per cell) configuration 130-10-10-18 Complete sci-FATE2 protocol will be available at protocols.io.

## Tn5 production and transposase assembly

Tn5 was produced as per [54, 61], to make 10*μ*M stock solution. This stock was diluted to 4*μ*M with dilution buffer (50 mM Tris, 100mM NaCl, 0.1 mM EDTA, 1 mM DTT and 50% glycerol), and transposase assembly was performed as per (Martin et al. 2022).

## Species mixture experiment

Mouse HM1 ES cells were cultured as per [37, 38, 54] in ES cell medium with 1,000U/ml LIF on feeder cells and human H9 cells in StemFlex on a laminin coating. Both were fixed in methanol + 0.1% DEPC as per above and stored at -80C. Cells were moved to ice for three minutes, then spun down at 2000rpm and 4°C for 5 minutes. Cells were resuspended in 1ml PBS + 0.1% DEPC + 3% v/v BSA, spun down again as before and resuspended in PBS at 1 million cells/ml. Mouse and human cells were then mixed in equal proportions and combinatorial indexing library preparation was performed as above.

## Cell loss quantification

Day 4 differentiation cells were collected and fixed in methanol as above. The modified chemical conversion protocol was carried out as described above, the original chemical conversion protocol was carried out as per [20]. Note that for this comparison, methanol fixed cells were used for both protocol, while the original protocol used cells fixed in 4% PFA. This was done to prevent differences in fixation from confounding differences in chemical conversion steps — however we and others have observed similar low yield of cells with original protocol and PFA-fixed cells [20]. For both, cells were initially counted and diluted to 5 million cells, and the number of cells remaining was estimated after all steps completed.

## sci-FATE pilot

The initial pilot of sci-FATE was performed with in vitro differentiating ES cells (day 4 of protocol), exactly as described in [20], with the following changes:1) no 4sU labelling was performed, ii) we ran 19 PCR cycles, iii) to concentrate the library, we collected 8 x 800*μ*l aliquots of library, and performed two rounds of Ampure bead concentration (0.7X then 0.8X) for a final library volume of 100*μ*l.

## RNA-seq and RT-qPCR

For both bulk RNA-seq and RT-qPCR, samples were collected and RNA isolated with a Qiagen RNeasy kit. For RNA-seq, samples were normalised to 375ng and library prep was performed with Kapa mRNA Hyperprep with polyA capture beads (KK8421). Libraries were sequenced using NovaSeq6000 with 25 million reads per sample. For RT-qPCR, 1*μ*g of RNA was reverse transcribed using Super-Script III first strand synthesis kit (Invitrogen) with random hexamers. PowerUp SYBR Green Master Mix (ThermoFisher) was used to perform qPCR with tenfold diluted cDNA, using Actin as a control.

## Martin et al protocol

For sci-RNA-seq2 comparisons, in vitro differentiating ES cells (day 4 of protocol) were fixed in methanol as described above. For library preparation, cells were washed once in 1ml PBS + 0.1% DEPC (where?) + 3% v/v BSA (NEB) then diluted to 1 million cells per ml in PBS and immediately submitted for library preparation. We performed combinatorial indexing protocol as above. For Martin et al 2022 protocol [60], after second-strand synthesis, we added 1*μ*l Qiagen protease per well and incubated for 30 minutes at 37C, then 20 minutes at 75C. We then performed tagmentation as above, before adding a quenching mix of 0.375*μ*l BSA (NEB), 0.375*μ*l 1% SDS and 2.25*μ* water to each well, incubating at 55C for 15 minutes. We then added 4*μ*l 5% Tween-20 to each well, proceeding to PCR with the addition of 2*μ*l primers and 20*μ*l NEBNext PCR mix. For the sample shown, we ran 16 PCR cycles, collected 3*μ*l per well, performed a 0.8X volume Ampure bead clean and isolated 200-600bp DNA by running the library through a 1% agarose gel and gel extraction (Qiagen), as performed in Martin et al 2022. In parallel, we ran an additional sample which we purified through a round of 0.8X Ampure bead cleaning followed by purification through a size-selection DNA column (Zymo) - we did not observe major differences in the data quality through this second approach.

## Raw data processing and label detection

Raw data processing and label detection was performed with the dynast package [22] and the GRCm39 reference genome. To estimate SNPs (and background conversion rate for estimation), dynast was run in ‘control’ mode, with a sci-RNA-seq2 sample from all time points of the differentiation mixed together, collected without 4sU labelling or chemical conversion. We found little benefit to using dynast’s consensus calling feature, so did not use it for analysis. We found that label rate estimation provided poor results, and so used dynast’s ‘count’ method instead (see supplementary note 1).

## Quality control and cell type classification

Basic processing was performed with ScanPy[62].To remove low quality cells, we set thresholds at the tenth and ninetieth percentiles for number of genes and UMIs observed per cell, and removed data that fell below or above these thresholds, respectively. This corresponded to an acceptable range of approximately 4000-25000 UMIs/cell and 2000-7000 genes per cell. We removed cells with less than tenth percentile labelling rate (approx. 10% labeled) and more than ninetieth percentile mitochondrial percentage of reads (approx. 5%). We used scrublet [63] to remove doublets with a doublet score threshold of 0.3. To handle batch effects, highly variable gene selection was stratified by replicate: 3000 highly variable genes were selected per replicate (excluding replicate 4 as an incomplete replicate), and only genes selected across all three replicates were retained (generally around 2000 genes). We found this successfully removed an evident batch effect between replicate 1 and replicates 2 and 3. For cell type classification, we adopted an ensemble approach: we clustered cells with the Leiden algorithm with high resolution (1.6), producing many more clusters than expected cell types (32 clusters, 9 expected cell types). We performed differential expression (using ScanPy) between clusters, scored cells based on key marker genes of expected cell types, and performed a basic classification based on binarised expression comparison to a knowledge table of marker genes, as per [2, 12]. For each cluster, we visualised the cluster in UMAP and MDE embeddings, visualised the keygene scores, the marker-based classification distributions and the top differentially expressed genes, and based on all analyses together, clusters were assigned to one of 10 groups: NMP, early neural, neural, pMN, MN, p3, V3, floor plate or other.

## Analysis of RNA velocity

For analysis of RNA velocity, we used scVelo to access RNA velocity datasets from human bone marrow, mouse dentate gyrus, mouse pancreas, and additionally accessed RNA velocity data from human and mouse neural tube[2, 11–13, 39, 40]. For visualisation of pancreas velocity data, we ran scVelo on default parameters, projecting velocities either using scVelo’s projection method, or by calculating directly projected velocity as:

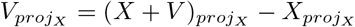

Where *proj*_*X*_ represents the application of *log*1*P* transformation followed by projection to PCA embedding of *X*.

## Comparison and quantitative benchmarking

To visualise velvet inference, we split the data into two: neural (early neural, neural, pMN, p3, floor plate, MN and V3) and NMP (NMP, mesoderm, early neural, neural and pMN from D3.2 to D5). Velvet was trained on default settings for both. As comparison, dynamo and scVelo were run on labelling and splicing data of the same systems respectively with default settings. For all three, visualisation was performed by predicting high dimensional velocities and directly projecting these velocities to PCA space.

For quantitative benchmarking, we split the dataset into six subsets: i) pMN/MN/FP, (approx. 10,000 cells) ii) p3/V3 (7,000 cells) iii) mesoderm (10,000 cells) iv) NMP system (18,000 cells) v) neural system (20,000 cells) vi) whole dataset (45,000 cells). These subsets were chosen to capture different resolutions of dynamics, from linear trajectories to systemic multifurcations. For each subset, scFates[41] was used to construct a pseudotime skeleton based on known biological trajectories. The parameters used for scFates were manually tweaked to ensure faithful recapitulation of known biological dynamics. With this skeleton, two methods were used to create a ground truth: 18 PTS: each cell was mapped to the closest position on the skeleton, and velocity was set as the displacement between that position and the subsequent position, ii) CRS: a pseudotime value for every cell was determined using sciFate’s pseudotime function, and this was passed to CellRank’s PseudotimeKernel[42], which was used to calculate a transition matrix which was then projected to PCA. For subset (vi) (whole dataset), an accurate pseudotime could be computed, but the underlying trajectory skeleton was not deemed to be reliable enough, so this dataset was not used for PTS assessment. Both ground truth velocities existed in 50D PCA space, and all tested velocities were also projected to 50D PCA space for comparison through cosine similarity. This projection was done directly, not using a nearest-neighbour projection heuristic. A third metric was used, cross-boundary direction (CBD), as done previously, using the implementation from [27]. With this method, data was clustered with Leiden algorithm, and directionality between specific clusters was defined. The agreement between inferred velocities and these directionalities was assessed by cosine similarity.

All comparison tools were run on default settings. Where a tool failed or crashed on a particular subset, it was assigned a score of zero (this only occured with velovi and smoothened data, occuring in 5/6 datasets in that instance).

## Velvet and svelvet velocity models

Velvet inference from metabolic labelling starts from the two equations:

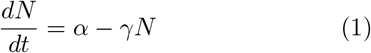

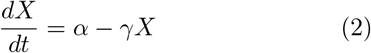

These represent the time derivatives for labelled (*N*) and total (*X*) reads, respectively, based on transcription (*α*) and degradation rates (*γ*). This framework assumes that the transcription rate for labelled and unlabelled reads is the same, for more details see supplementary note 1. Since we can assume no initial conditions, we can solve the labelled derivative equation to give:

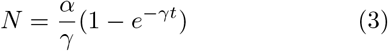

This can be rearranged to give an equation for transcription rate:

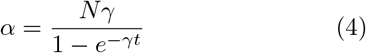

Inserting this into the equation for total velocity, we get our key equation:

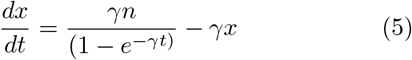

This defines velocity in terms of three observables (new reads, n; total reads, x; labelling time, t) and one latent parameter, the degradation rate, *γ*. Unlike RNA velocity, there is no need to infer splicing parameters, initial conditions or gene-specific latent times, resulting in a simpler modelling framework. This equation can be rearranged to provide a prediction of labelled/new reads based on predicted total expression and velocity:

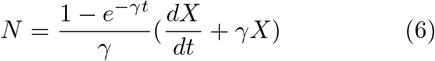

This equation is built into the loss function of velvet.

For svelvet, we start with a similar framework:

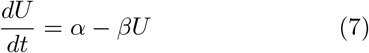

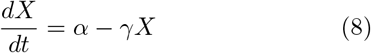

We assume *β* = 1 as per previous work. At steady-state (when ^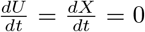^, these equations can be equated to give 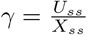, providing a velocity definition:

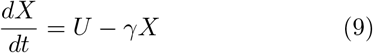

Which, as with velvet, is reframed to predict unspliced reads from predicted total expression and velocity. This is distinct from most previous velocity approaches which model unspliced-spliced ratio. This is done to bound *γ* between zero and one, with the aim of improving stability in the face of highly variable unspliced proportions across genes (Figure 1A).

To train either velvet or svelvet, we input new and total datasets, along with precomputed neighborhood indices for the neighbourhood constraint. Optionally, we can include a precomputed transition matrix calculated based only on distances (as per [64]), to balance cell similarity with velocity direction when producing the velocity projection as part of the neighbourhood constraint - following ideas developed in [11, 43] (however, initial exploration found that doing so had little effect on results).

The above framework is built into a variational autoencoder (VAE) framework [34]. Briefly, the VAE is composed of two neural networks, an encoder and a decoder. The encoder maps the input data to a lower-dimensional ‘latent’ space. This mapping is probabilistic, meaning that the encoder learns to map to mean and variance parameters of latent variables, assumed to be independent Gaussians, and a latent representation of the data is sampled from these learned Gaussian distributions. The decoder reconstructs the original data from this sampled latent representation. The training involves optimizing the parameters of the encoder and decoder networks through a variant of stochastic gradient descent, where the objective is to minimize the reconstruction loss (the difference between the input and the reconstructed data) and a regularization term, the Kullback-Leibler (KL) divergence, that forces the learned distribution to be close to a standard normal distribution. This dual-objective function effectively results in a balance between data fidelity and statistical regularization.

In velvet, the latent representation of the data is passed through a latent vector field (a third neural network) that produces velocity predictions. These latent velocities are also projected through the decoder to produce reconstructed high dimensional velocities, which along with the data reconstruction and equation 6, is used to create a reconstruction of new (or labelled) reads. Thus an additional term is added to the loss function, comparing the original to reconstructed labelled read dataset (for this, we calculate log-mean-squared error). The neighbourhood constrain forms an additional component of the loss function.

The model is trained in two steps: first, the VAE and the vector field are trained together, then the VAE parameters are frozen and the vector field is trained with the neighborhood constraint. By default, the first stage is performed for 200 epochs and the second for a further 800. We use the entire dataset as one batch, and use a learning rate of 0.001, and the AdamW optimiser with weight decay of 0.001. By default, we use 50 latent dimensions, a linear decoder, and perform the neighborhood constraint in latent space.

To initially approximate *γ* with labelling data, we start with the assumption that steady-state dynamics are observed, where *N* = *kX*_*ss*_ and 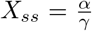, where k is a proportional value representing the proportion of total that is labelled at steady state. Plugging these into equation 4 and rearranging for *γ* gives us:

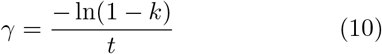

Thus, we perform extreme-value regression of new and total reads as per previous studies [25, 43], and calculate *γ* from the estimated slope from these regressions, *k*.

For splicing data, we cannot define an equivalent of equation 4 without initial conditions or splicing parameters, so we use a more simplistic approximation of *γ*, performing unspliced-total extreme-value regression, and setting *γ* = *k*.

## Neighbourhood constraint

To create projections of predicted velocities, we implemented a neighborhood based projection method as described previously [43], but instead of projecting into a new embedding, we use the method to ‘project’ into the same space, thus allowing comparison between the original velocities and projections that are constrained by the distribution of local neighbour cells. The projection method in brief: we calculate the cosine similarity between a cell’s velocity vector and the displacement between the cell and all its nearest neighbours (default 100 neighbours). We apply a softmax transformation to convert these scores to transition probabilities (applying a kernel width parameter *σ*^2^, default 0.1). We then calculate the velocity vector’s expected displacement with respect to the cell’s transition matrix, adjusting for non-uniform data density as per [43].

We compare the neighbourhood-based expected displacement with the original velocity vector with a cosine embedding loss (1*−*cos(*V, V*_*proj*_)) where cos is cosine similarity. This component of the loss is combined with the vector field loss comparing observed and reconstructed labelled data, in the second phase of training. The two only differ when the projection fails to accurately reconstruct the original vector, which will occur when predicted velocity vector points away from neighbours. In which case, the neighbourhood constraint will encourage predictions to point towards local neighbourhoods. Cell neighbour indices are precomputed before training, and neighbourhood projection is performed in a broadcasted fashion, allowing a computationally efficient implementation of the neighbourhood constraint.

## VelvetSDE dynamical model

To train velvetSDE, we embed the trained velvet vector field as the drift component of a neural stochastic differential equation system[35, 36] of the following form:

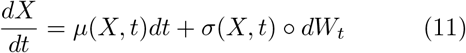

Where *μ*(*X, t*) is a neural vector field, *σ*(*X, t*) is a noise function and *W*_*t*_ represents a Weiner process of Brownian motion. To reduce memory usage, adjoint-mode gradient computation is used. The SDE is constructed as a Stratonovich integral solved using the midpoint method. This is done as Stratonovich integrals have lower computational cost for adjoint methods than Ito integrals [35]. For this study, *σ*(*X, t*) = *k*, where *k* is a user-defined scalar.

In parallel, we define a Markov process model from the learnt vector field: we calculate a transition matrix, *π*, from each cell *i* to its neighbours:

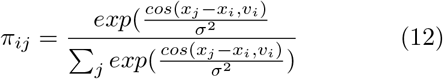

As per the transition matrix calculation for neighbourhood constraints. We can define a Markov process random walk as:

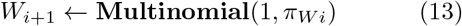

In words: The next state is chosen as a sample from a multinomial distribution with a probability vector equal to the current state’s transition probabilities, defined by *π*. We thus iteratively define random paths through the data manifold determined by the velocity-guided transition matrix. We can optionally define terminal states (where *W*_*i*+1_ = *W*_*i*_) and can produce a cubic spline of the Markov path so that the number of timesteps can equal that of nSDE simulations, without requiring that the same number of Markov steps and nSDE steps are taken.

To train velvetSDE, for each cell in a batch, we simulate *n* trajectories with the nSDE model (*X*^*SDE*^) and *n* trajectories with the Markov process random walk (*X*^*MRW*^). Each cell’s loss is defined as the average of the Kullback-Leibler divergence of the SDE simulation from the Markov random walk, stratified by timestep:

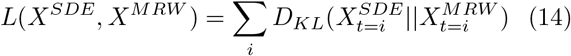

In words, for each timestep, this is the Kullback-Leibler divergence of that timestep’s distribution of nSDE points from that timestep’s distribution of Markov simulations, assuming both distributions to follow a Gaussian distribution. This measures how far from the Markov simulation the nSDE simulation has moved. For training, we set the noise magnitude of the nSDE to roughly equate the noise magnitude of the Markov random walk, in this study, a noise magnitude of 0.2 is used for this. We train for 250 epochs, running 200 cells per epoch and 50 simulations per cell (each, for nSDE and Markov random walk), with same learning rate and optimiser as above.

## Decision boundary analysis

To cluster trajectories, we performed k-means clustering on trajectories, reshaped to be (*n, t ×d*) where *n* is the number of trajectories, *t* is the number of timesteps and *d* is the number of latent dimensions. To define decision boundaries, we simulate each cell’s trajectory ten times, cluster all trajectories, and define ‘mixed-fate’ cells within a decision boundary as any cell with trajectories belonging to multiple clusters. To investigate the role of noise in the temporal evolution of decision boundaries, we performed neighbourhood smoothing of timepoints to create a continuous scale of time, binned cells into time windows, and performed the above decision boundary analysis for each time window across multiple levels of noise magnitude in velvetSDE’s diffusion component. Time smoothing was performed to accommodate the uneven distribution of timepoints (D3.2, D3.4, D3.6, D3.8, D4 and D5), and analysis provides a proof of principle, though the specific timings may be biased by this uneven sampling. For in silico perturbation, we took ‘mixed-fate’ cells, and set each TF’s expression value in turn to the maximal observed value for that TF, we then projected these perturbed data to the velvetSDE latent space, and simulated each cell’s trajectory. We classified trajectories by assigning each trajectory to the trajectory cluster (i.e. neural or mesoderm) that it is on average closest to. To determine negative log p-values, we performed binomial tests, where *k*=observed number of neural trajectories, *n*=number of simulated trajectories and *p*=observed wildtype probability of neural trajectories from mixed-fate cells. Note that we tested in silico ‘knock-outs’ as well as ‘over-expressions’ i.e. setting gene expression to zero, and found the expected inverse results (as expected) however trends were less clear, possibly due to the confounding factor of drop-out being modelled in Velvet’s zero-inflated negative binomial fitting. In other words, zero expression is caused by lack of expression and by dropout; maximal expression is only caused by maximal expression.

## Gene expression analysis

To produce gene expression distributions from trajectories, we project them through the model’s decoder to gene expression space, taking the decoded mean value as the expression level. To produce orbital phase portraits, we scaled the distribution of values each gene between 0 and 1 and performed Gaussian kernel smoothing with *σ* = 5. To produce trajectories to analyse, we simulated all D4 cell trajectories. All trajectories were done for 100 steps. For Markov random walks, we simulated 50 steps with a transition matrix constructed using 10 nearest neighbours. We used cubic splines to interpolate 50 steps into 100 timepoints. For pseudotime binning, we subsetted cells early neural, neural pMN and MN cells from day 4 and day 5. We calculated velocity pseudotime using scVelo, and sorted cells into 100 bins based on pseudotime score. For each simulation, cells were randomly chosen from each bin and trajectories constructed as the path across pseudotime bins. To score trajectories on conservation of expected gene orderings, we used seven orderings: *Sox*2*→ Olig*2, *→ Irx*3*→ Olig*2, *→ Pax*6 *→ Olig*2, *→Olig*2 *→ Neurog*2, *Neurog*2*→ Mnx*1, *→ Neurod*4*→ Isl*1, and *Mnx*1 *→Tubb*3. To assess ordering, we compared the argmax of each gene being compared for a given trajectory. If a trajectory contained all orderings correct, it scored 1, otherwise 0, and the score reported is the mean across all trajectories for a given method.

## Neural patterning analysis

Trajectories were averaged by taking the median of each timestep across clustered trajectory distributions. To assess the temporal progress of fate probabilities, trajectories for cells at each timepoint were simulated, and simulations were compared to average trajectories constructed from D4. Dynamic time warping was used in this distance measurement to account for the variable start point of trajectories across timepoints.

## Computational resources for analysis

All analysis was run on a cluster, accessing a single GPU core from NVIDIA V100 GPU node and 8 CPU cores, with 120Gb of memory. In this configuration, for approx. 20,000 cells, velvet trains within five minutes, velvetSDE within one minute.

## Code and data availability

All code for raw data processing and label detection with dynast is available at: https://github.com/rorymaizels/sciFATE2_processing. All code for computational analysis is available at: https://github.com/rorymaizels/Maizels2023aa. The deep learning framework, available as the software package, velvet, can be installed from: https://github.com/rorymaizels/velvet. Raw data and processed data can be accessed with the GEO accession: GSE236520.

## SUPPLEMENTARY MATERIALS

### Supplementary Note 1

Previous work has found that 4sU incorporation rate is roughly 1 in 40 uridines [23], implying that a significant proportion of nascent reads would not possess a detectable 4sU label within the sequenced read. This has previously been resolved with a Bayesian approach [22, 65] to estimate the experimental T-to-C conversion rate and infer the new-total ratio for each gene in each cell with a binomial mixture model. This Bayesian approach is implemented in dynast with two modes: ‘*π*_*g*_ mode’, which implements the original approach to infer a parameter for each gene in each cell, or *α* mode, which instead infers a cell-wise correction value by which the number of labelled reads is adjusted across all genes for a cell. The motivation for the *α* mode is to make the problem more tractable for the sparse, noisy data of single-cell sequencing. We found that dynast’s *π*_*g*_ implementation failed even basic tests (such as displaying a positive correlation between new and total reads), and so we progressed with *α* mode instead. With this method, we observed an increase in the proportion of reads labelled as new from 20% to 40%, consistent with the expectation that as many as half of new reads would not contain a detectable label (Figure S4A). However, visualising new vs. old reads for estimation and counting methods for a given gene (Olig2, for example, in Figure S4B) showed a large increase in the number of cells predicted to possess only ‘new’ reads, and the proportion of cells that possess both new and old reads (and thus, contain useful dynamical information) is consistently reduced with the estimation method versus counting (example shown for Olig2, Figure S4C). We also found that across all genes, with estimation there was a consistent positive correlation between the total number of reads detected in a cell and the label rate detected in a cell, which was absent when using the counting method. We found that steady-state degradation rate approximations from counting data gave a half-life distribution closer to the experimentally measured half life distribution of mRNAs in mES cells, when compared to degradation rates inferred from estimated labelling. Finally, testing the dynamical information with estimation vs. counting with velvet and dynamo, we found that using estimated label rates substantially worsened velocity predictions (Figure S4F).

Based on these findings, we chose not to use the Bayesian estimation method for our analysis, finding that inference of dynamics generally performs well. Our velocity model is defined in terms of observed ‘new’ and ‘total’ reads, along with a degradation parameter that is itself initially estimated from the observed ‘new’ reads. As such, velocity is defined relative to the observed ‘new’, and the effect may be only that the magnitude of velocity vectors may be underestimated.

Moreover, our results are not consistent with the observation that as many as half of nascent reads may be undetectably labelled, and it may be that recorded 4sU incorporation rates are underestimations: these figures come from studies that look at incorporation rates in bulk RNA[66](which would include many species such as rRNA with considerably longer half lives than mRNA) or with below-saturation levels of labelling[67] (e.g. 100*μ*M) for labelling periods of only 1-3 median mRNA half lives (12-24 hours). As a result, 4sU may not have fully saturated the transcriptome in these studies, leading to an underestimation of incorporation rate. More work on the measurement and Bayesian inference of 4sU incorporation will be needed to explicate this matter.

**Supplementary Figure 1:**
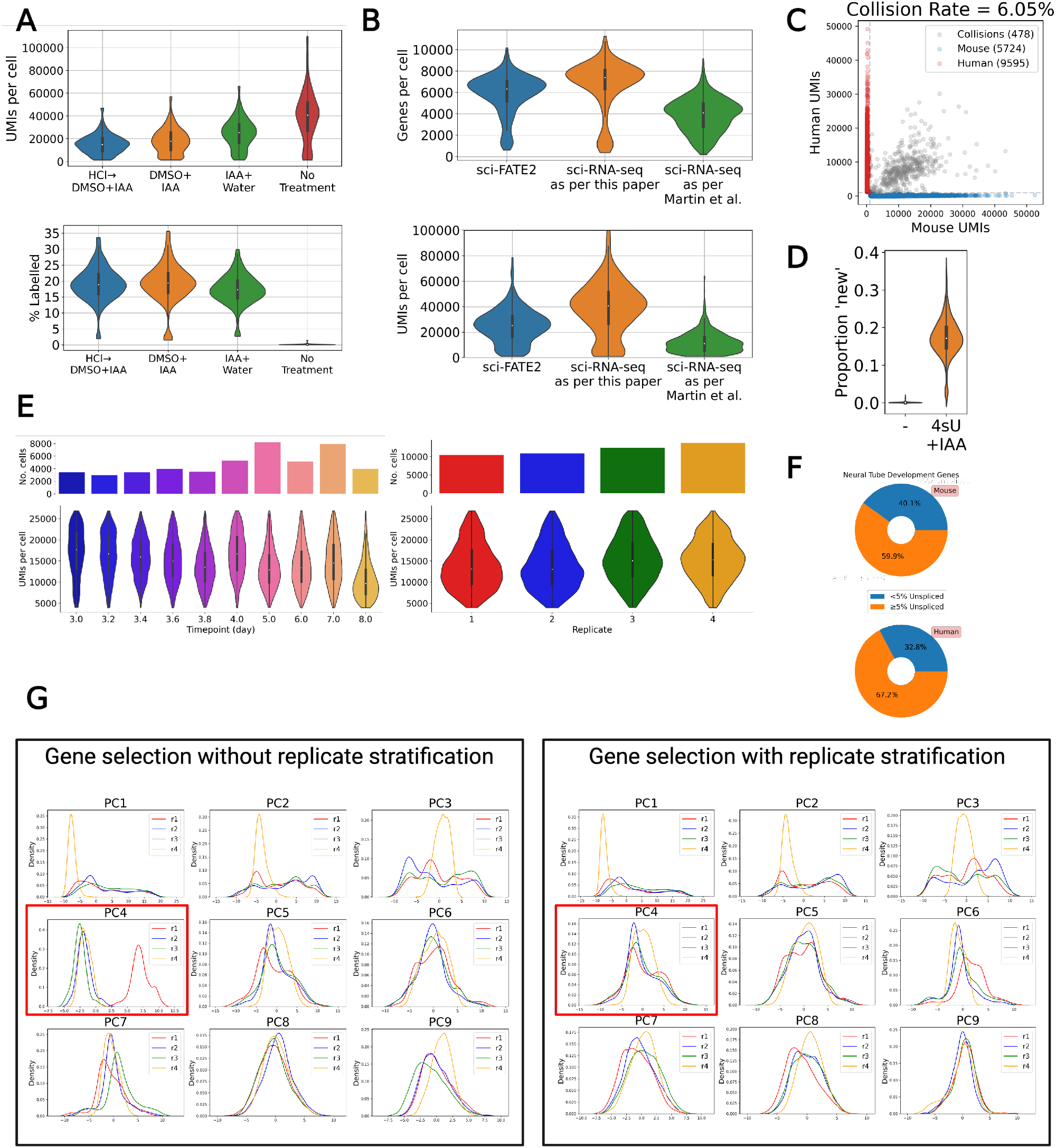
Data Processing and Quality Control. A. Comparison of UMis per cell and label rate for different chemical conversion protocols, B. Comparison of UMIs and genes per cell for the combinatorial indexing protocol used in this study compared to the protocol in Martin et al. 2022. Protocols were run in a single experiment, sequenced together and demultiplexed afterwards. C. Species mixture experiment measurement of doublet rate. D. Comparison of detected ‘labelled’ reads with dynast, with and without 4sU + IAA. E. UMIs per cell distributions and cells per condition, shown for timepoints and replicates. F. Proportion of 177 key neural tube development markers [2] that have less than 5% unspliced reads in human and mouse. G Distribution of principal of data, split by replicate, shown for gene selection without (left) and with (right) stratifying selection by replicate, emphasising the batch effect observed in and removed from component 4.

**Supplementary Figure 2:**
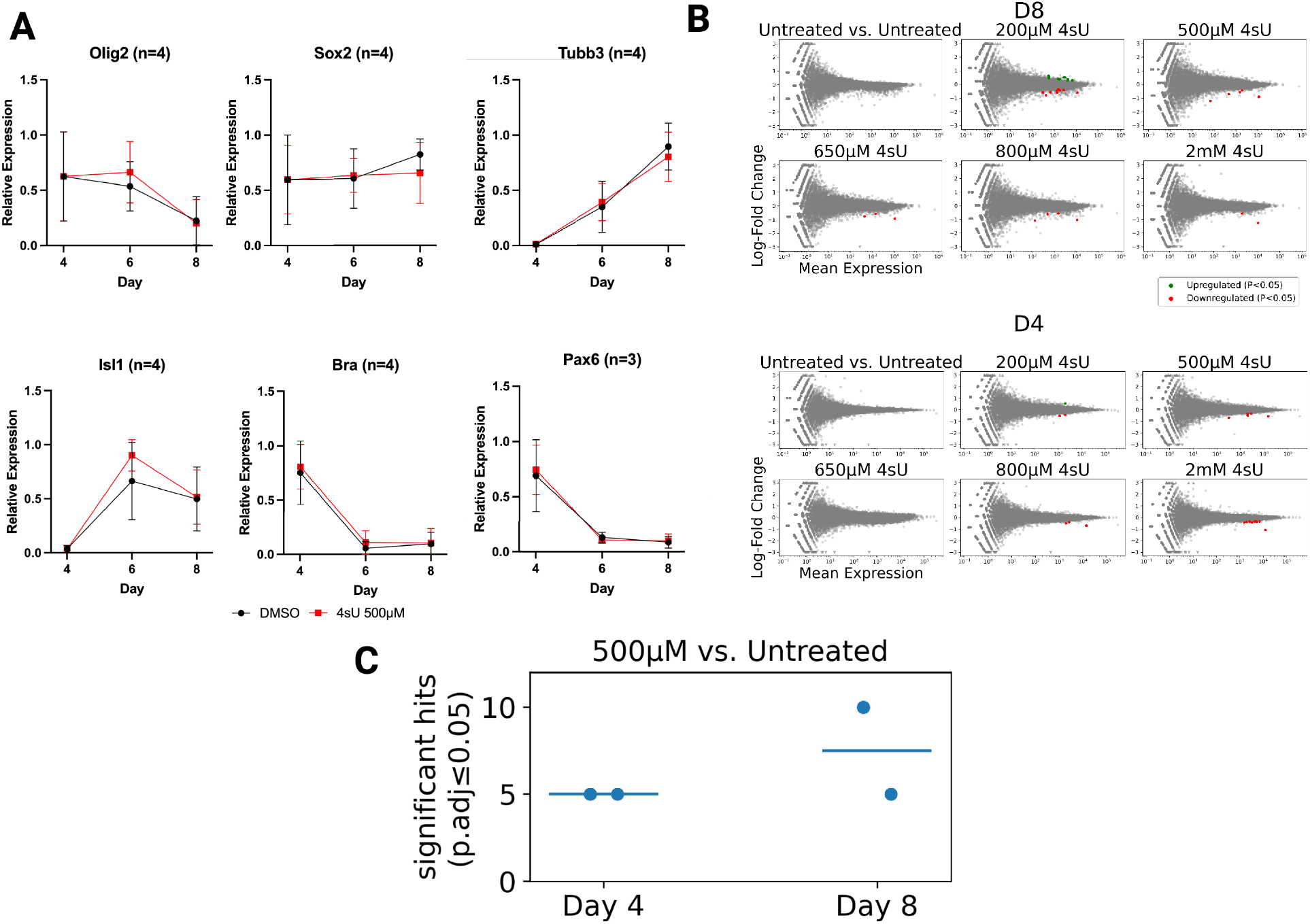
4sU Labelling QC. A. RT-qPCR of key genes with and without 2 hour 500*μ*M 4sU labelling before collection. B. RNA-seq of samples with and without 2 hours of varying concentrations of 4sU labelling before collection on day 4 and 8 of differentiation. C. Summary of RNA-seq results for 500*μ*M 4sU for two hours, across two replicates.

**Supplementary Figure 3:**
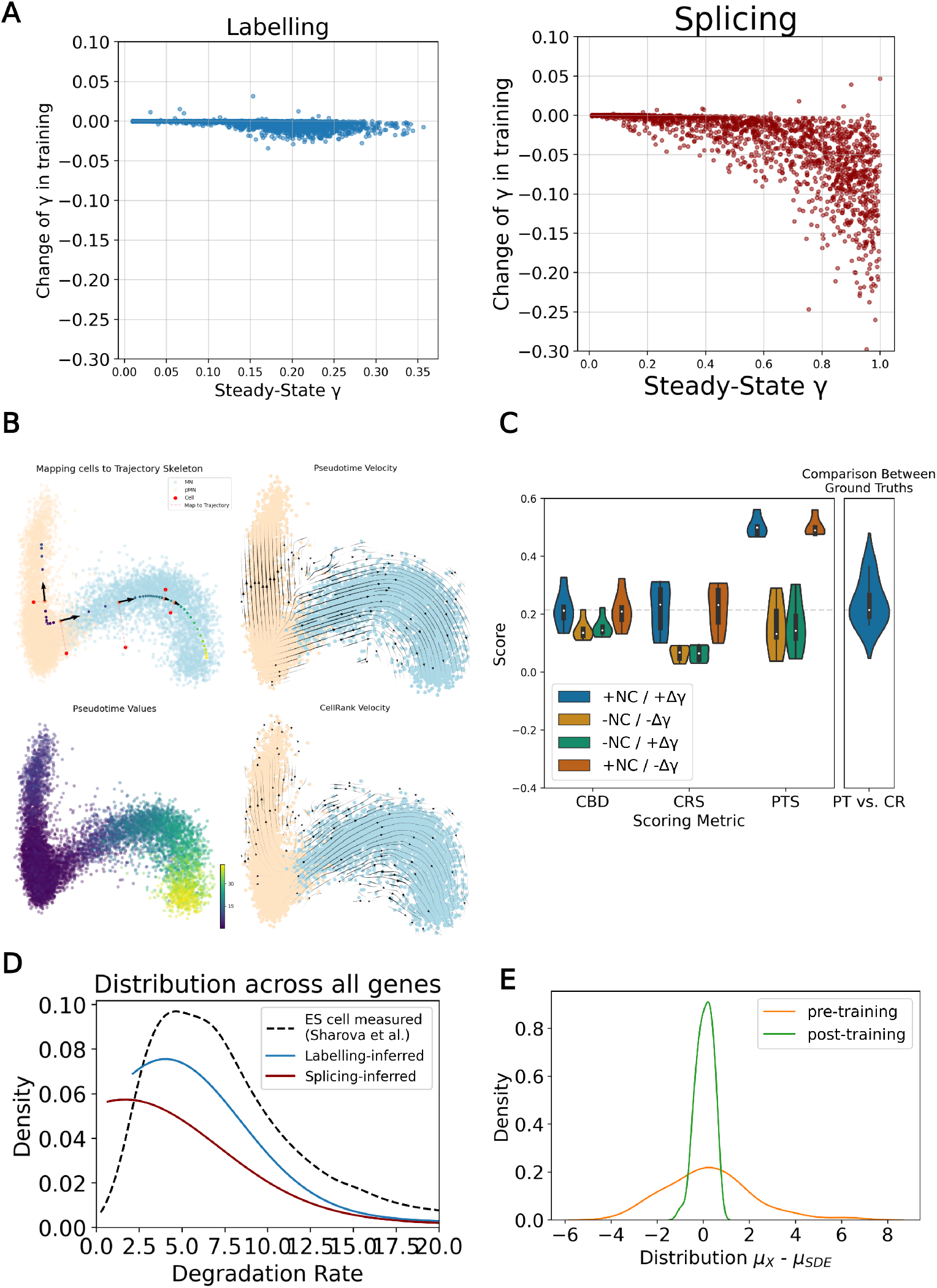
Velocity inference. A. Change in gamma parameter through training for labelling and splicing data. B. Comparison of quantitative benchmarking scores for velvet with and without modelling components, NC: neighbourhood constraint, Δ*γ*: learnable degradation parameter. C. Benchmark ground truth construction overview; a pseudotime skeleton was constructed with scFates (top left); cells were mapped to this skeleton and velocities set to closest point’s displacement to next point for pseudotime score (PTS; top right); pseudotime values were constructed with scFates (bottom left) and CellRank PseudotimeKernel was used to construct an alternative velocity ground truth for CellRank score (CRS) (bottom right). D. Comparison of distribution of gamma values inferred from splicing and labelling data, compared to experimentally measured values in ES cells from [68] E. Across latent dimensions, the distribution of data mean minus simulation mean for pre- and post-training velvetSDE simulations.

**Supplementary Figure 4:**
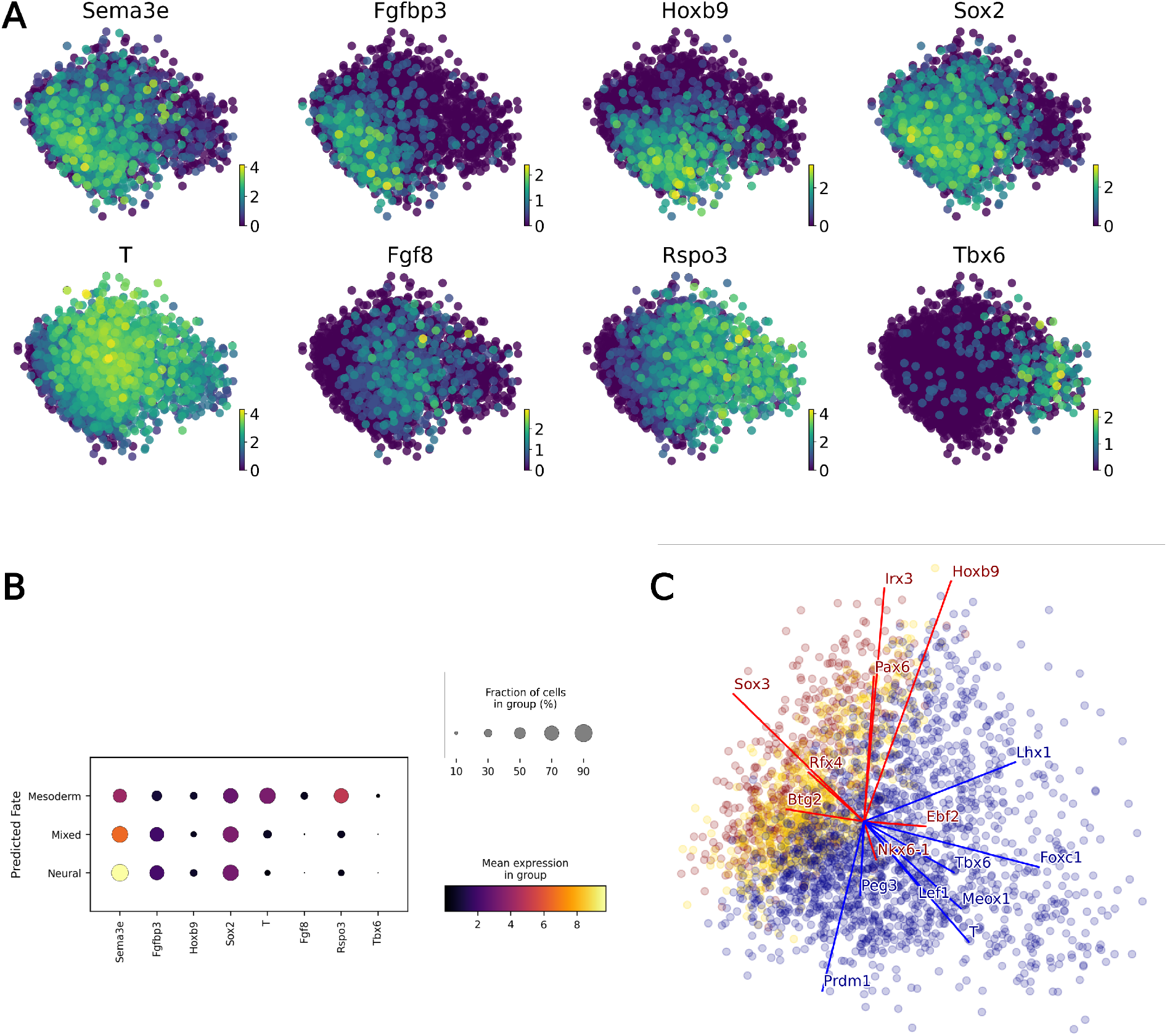
NMP fate prediction. A. Expression of key markers on D3, prior to RA addition. B. Expression of key markers across cells predicted to have neural, mixed or mesoderm fates, for D3.2 cells. C. Gene loading of top in silico perturbation hits in PCA of latent space (with linear decoder, gene loading vectors can be visualised similarly to PCA).

**Supplementary Figure 5:**
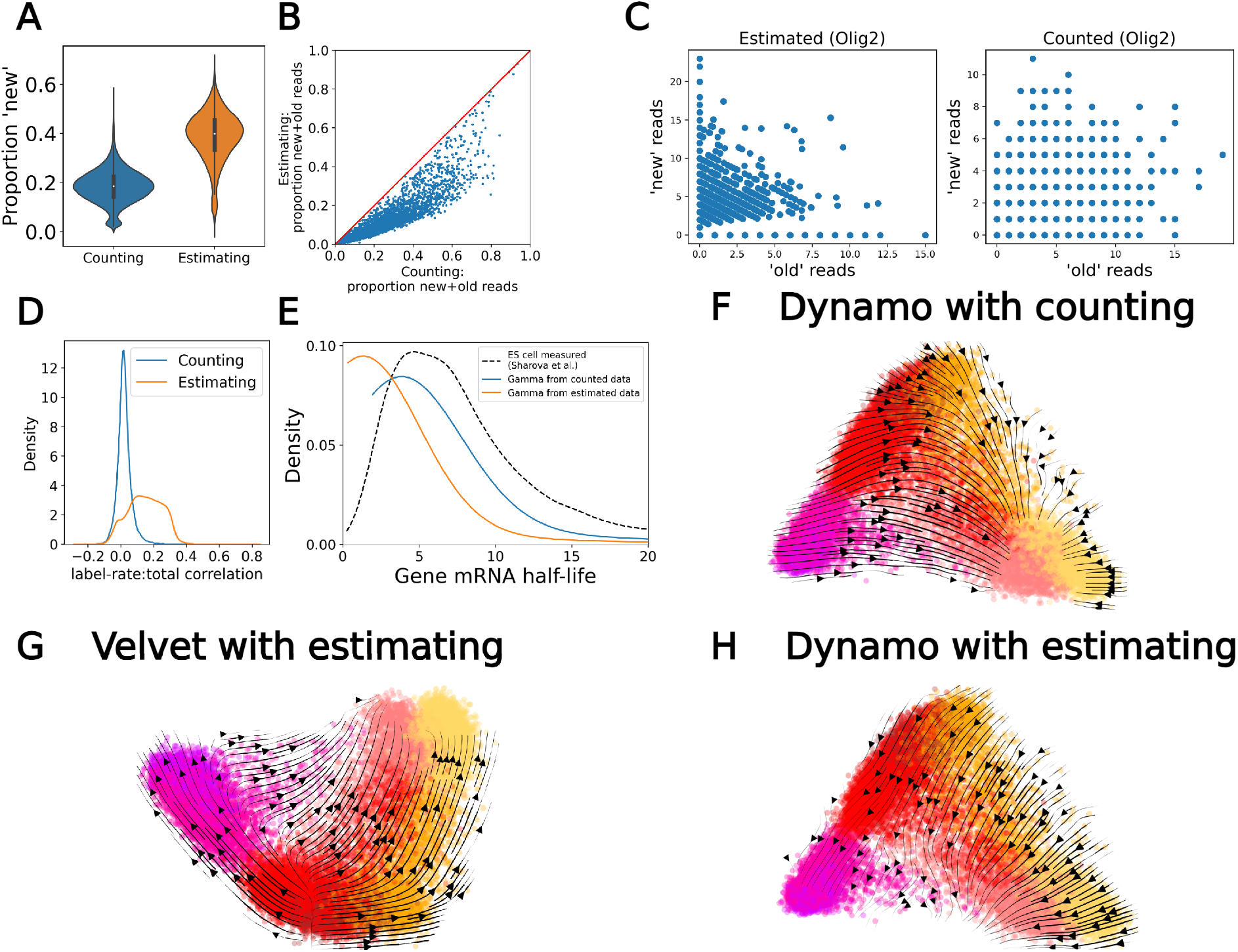
Estimated vs. counted labelled reads with dynast. A. Proportion of reads classified as ‘new’ (or ‘labelled’) with counting or estimating methods. B. Scatter of new and old reads across cells for Olig2, comparing estimating (left) and counting (right) methods, C. Across genes, comparing the proportion of cells that contain both new and old reads together for counting and estimating methods, D. Distribution of correlations between label rate and total counts across genes for estimating (orange) and counting (blue) methods. E. Distribution of predicted gammas using counted (orange) and estimated (blue) methods, compared to experimentally measured distribution in mES cells, F. Velocity dynamics for Dynamo with counting, G. Velvet with estimating data. H. Dynamo with estimating data.

**Supplementary Figure 6:**
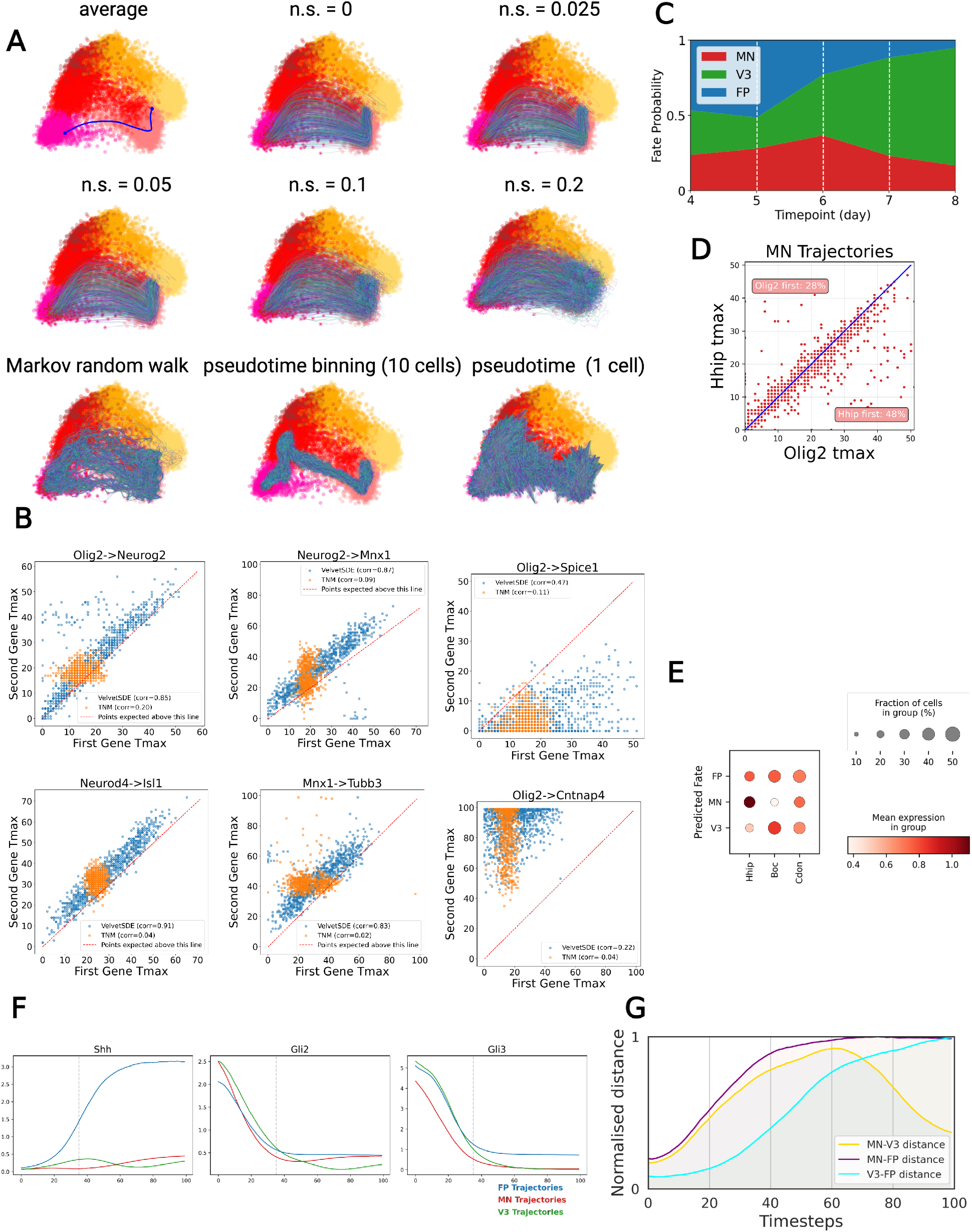
Neural system dynamics. A. Latent space PCA visualisation of trajectories used for gene ordering scoring. B. Comparison of expression timings for velvetSDE simulations (noise = 0.1) and technical noise model. C Streamplot of Predicted fate probabilities across timepoints. D. Coexpression of Hhip and Olig2 in motor neuron trajectories visualised. E. Expression profile of Shh signalling genes in cells of different predicted fates. F. Median expression profile for Shh and Gli proteins across trajectories. G. Normalised distances between median trajectories of each fate.

**Supplementary Figure 7:**
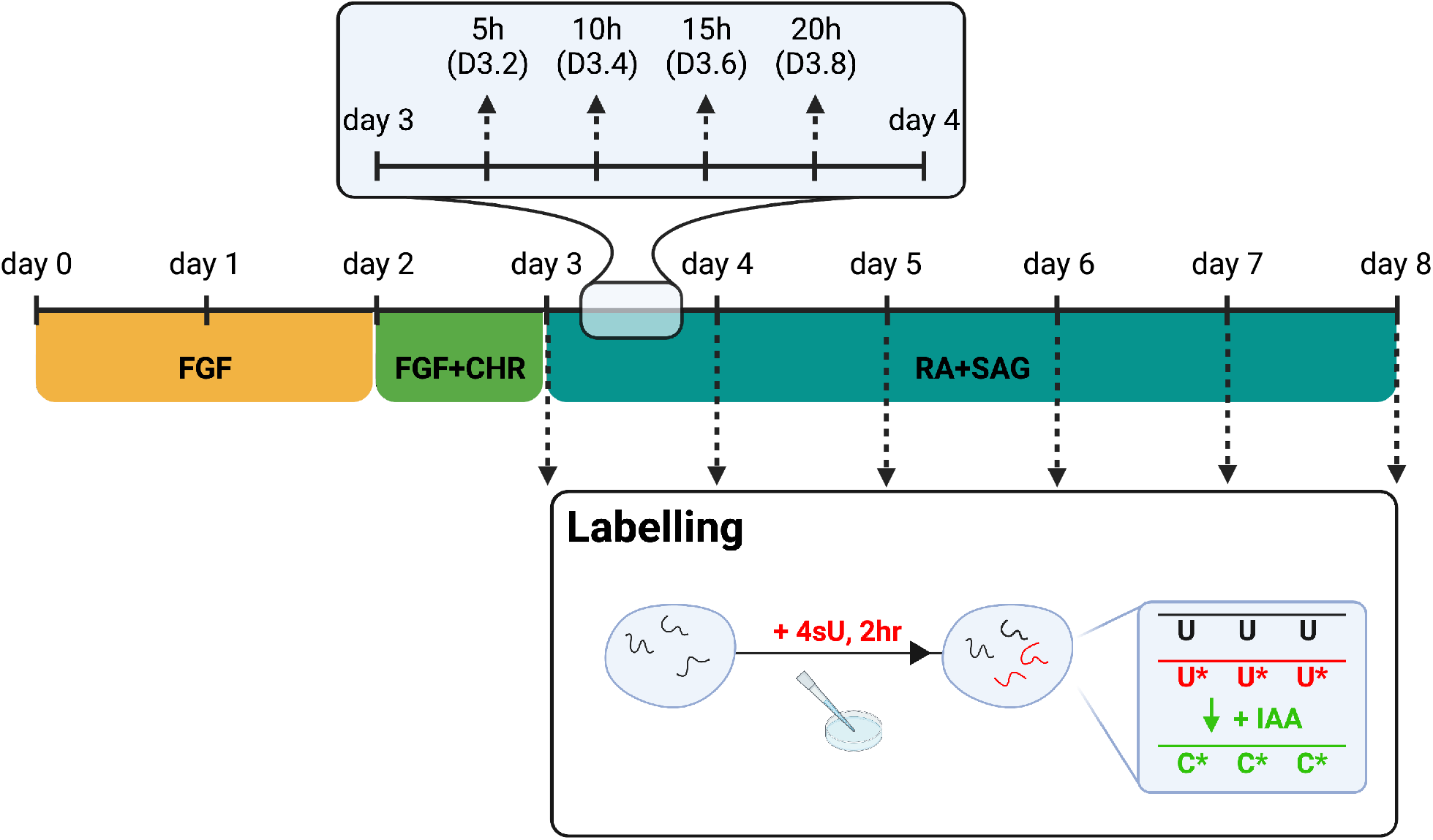
In vitro neural differentiation protocol. Schematic outlining in vitro protocol with collection points and 4sU labelling regime.

